# Mathematical modelling of brain mTOR activity identifies selective vulnerability of cell types and signalling pathways

**DOI:** 10.1101/2022.12.15.505685

**Authors:** Alexandra Pokhilko, Hannah Sleven, M. Zameel Cader

**Affiliations:** Translational Molecular Neuroscience Group, Nuffield Department of Clinical Neurosciences, University of Oxford, Oxford, OX3 9DS, United Kingdom

**Keywords:** mTOR, mathematical modelling, ordinary differential equations, RNA sequencing, cerebrovascular disease

## Abstract

The mTOR pathway is a global regulator of protein biosynthesis and cellular homeostasis. Understanding the differences in mTOR pathway activity between cell types is important for elucidating the role of mTOR in physiological and pathophysiological processes. The non-linear structure of the pathway, with multiple feedback loops and inputs complicates the interpretation of experimental data and requires mathematical modelling. We modelled mTOR activation under healthy and disease conditions using recently published single nuclei gene expression data from the human brain. The model predicts substantial variations in mTOR pathway activity between cell types, with neurons and astrocytes being highly sensitive to insulin and neuregulins, while vascular smooth muscle cells and pericytes are highly sensitive to PDGF. Two principal negative regulators of mTOR - TSC and PTEN have prominent roles in PDGF-mediated mTOR activation in endothelial cells and oligodendrocytes. In Alzheimer’s Disease brain we find that insulin-mediated activitation of mTOR pathway is selectively upregulated in microglia and oligodendrocytes and downregulated in other cells. Our mathematical modelling characterises ligand sensitivity of cerebrovascular cell types and provides insights into the brain mTOR dynamics in health and disease.

## 1. Background

The mTOR kinase pathway controls cellular growth by adjusting protein biosynthesis to external and internal conditions, including nutrient and growth factor availability (1, 2). In healthy mature brain tissues, the canonical mTOR pathway is activated by secreted signalling ligands (3, 4). This activates a signalling cascade through tyrosine kinases, leading to phosphorylation and activation of mTOR pathway components, such as PKB (AKT), mTORC1/mTORC2 complexes and S6 kinase (S6K). Activated S6K is one of the main effectors of the mTOR pathway, accelerating protein translation by phosphorylating ribosomal S6 protein (1–3).

Brain tissue is a complex mixture of multiple cell types, each with their own unique metabolic demands and functions. Given the central role of the mTOR pathway in regulating cell metabolism, cell-type specific variation in mTOR pathway activity and responsiveness provides a potential means by which each cell type can thrive in the complex brain environment. Simplistically, control of mTOR responses could be dependent on the expression of receptors for ligands, such as insulin and growth factors. However, variation in the expression of the many components of the mTOR pathway offers additional and complementary cell-internal regulatory mechanisms. The mTOR pathway includes multiple feedback loops, further complicating the analysis of mTOR responses to perturbations (5). The impact of the pathway complexity is evidenced in previous efforts to restrict growth of cancer cells by inhibiting mTOR, which often had the opposite result - upregulation of tumour growth through a relief of negative feedback loops (5). This complexity makes mathematical modelling of the mTOR pathway a necessary tool in the planning and interpretation of mTOR-related experiments (1). However, existing models do not address the differences in abundance of mTOR pathway components between cell types. This so far precluded quantitative analysis of mTOR dynamics in different types of brain cells.

Our analysis of publicly available high resolution single nuclear RNA-seq (snRNA-seq) data (6) indicated that human brain cells have significantly different levels of expression of mTOR pathway components, suggesting potential cell type specific responses to physiological signals. To explore the effects of variations in the levels of mTOR components on activity of mTOR pathway, we built a mathematical model of the canonical mTOR pathway, operating in the brain cells. This revealed substantial differences in mTOR responses to stimulation between cell types, and identified cell type-specific dependency of mTOR pathway activity on the levels of tyrosine kinase receptors and other components of the mTOR pathway. Our analysis also highlighted cell-type and ligand specific changes in mTOR pathway activity under pathological conditions.

## 2. Results

### 2.1. A mathematical model of mTOR activation

We first constructed a reference model of the canonical mTOR pathway (Supplementary Information; Figs. S1,S2) based on integration of two previously published models (7, 8) of insulin and PDGF signalling. We chose these two models because they were constructed based on detailed experimental data on individual components of mTOR pathway. In addition, the models were relatively simple, but catching key observations, e.g., presence of a negative feedback regulating insulin receptor and sigmoidal dependence of mTOR pathway activity on a wide range of stimuli. From the recently published human brain snRNA-seq data (6) we analysed growth factor (GF) and insulin receptor expression in previously classified 13 cerebrovascular cells types: 3 endothelial cells subtypes, 2 pericytic subtypes, 2 smooth muscle cell subtypes, 2 astrocyte subtypes, microglia, oligodendrocytes, oligodendrocyte precursor cells and neurons. We identified three main groups of ligands and their receptors, that are expressed in the brain cells and participate in the activation of mTOR via the canonical pathway (3, 4). The relevant receptors included PDGF receptors (PDGFR); neuregulin (NRG) receptors (NRGR) of the EGF family; and insulin receptors (IR) to insulin-like ligands (insulin/IGF; Supplementary Information; Fig. S3). We constructed the model to be able to analyse the responses to these three groups of ligands with model parameters fitted to published data on the kinetics of mTOR activation by insulin, PDGF and NRG (7–9), as well as the data from TSC knock out (TSC-KO) and rapamycin treated cells (Supplementary Information; Figs. S1,S2; Table S1).

The model is represented by a system of ordinary differential equations (ODE), describing mTOR activation by insulin-like and non-insulin types of ligands (Fig. 1). The activation of mTOR by these two groups of ligands differs in the initiation steps. Thus, insulin-like ligands bind to pre-assembled cross-linked dimers of insulin receptors (IR), with only one molecule of a ligand bound to the receptor (10). The non-insulin ligands (e.g. PDGF) form monomeric complexes with their receptors, following by their dimerization (8). In addition, mTOR activation by insulin-like ligands includes specific steps, such as phosphorylation of insulin receptor substrate (IRS1), as well as feedback loops absent from the PDGF activation pathway. The insulin and non-insulin pathways have common steps, including Tyr and Ser phosphorylation of PKB and activation of mTORC1, mTORC2 and S6K (Fig. 1). The published models were modified to integrate them together; incorporate negative feedback through inactivation of IRS1 by mTOR; and add the key inhibitors PTEN and TSC (Supplementary Information). The input to the model is the step-wise addition of different concentrations of ligands for 2 hours and the output is the resulting levels of S6K activation.

**Figure 1.**
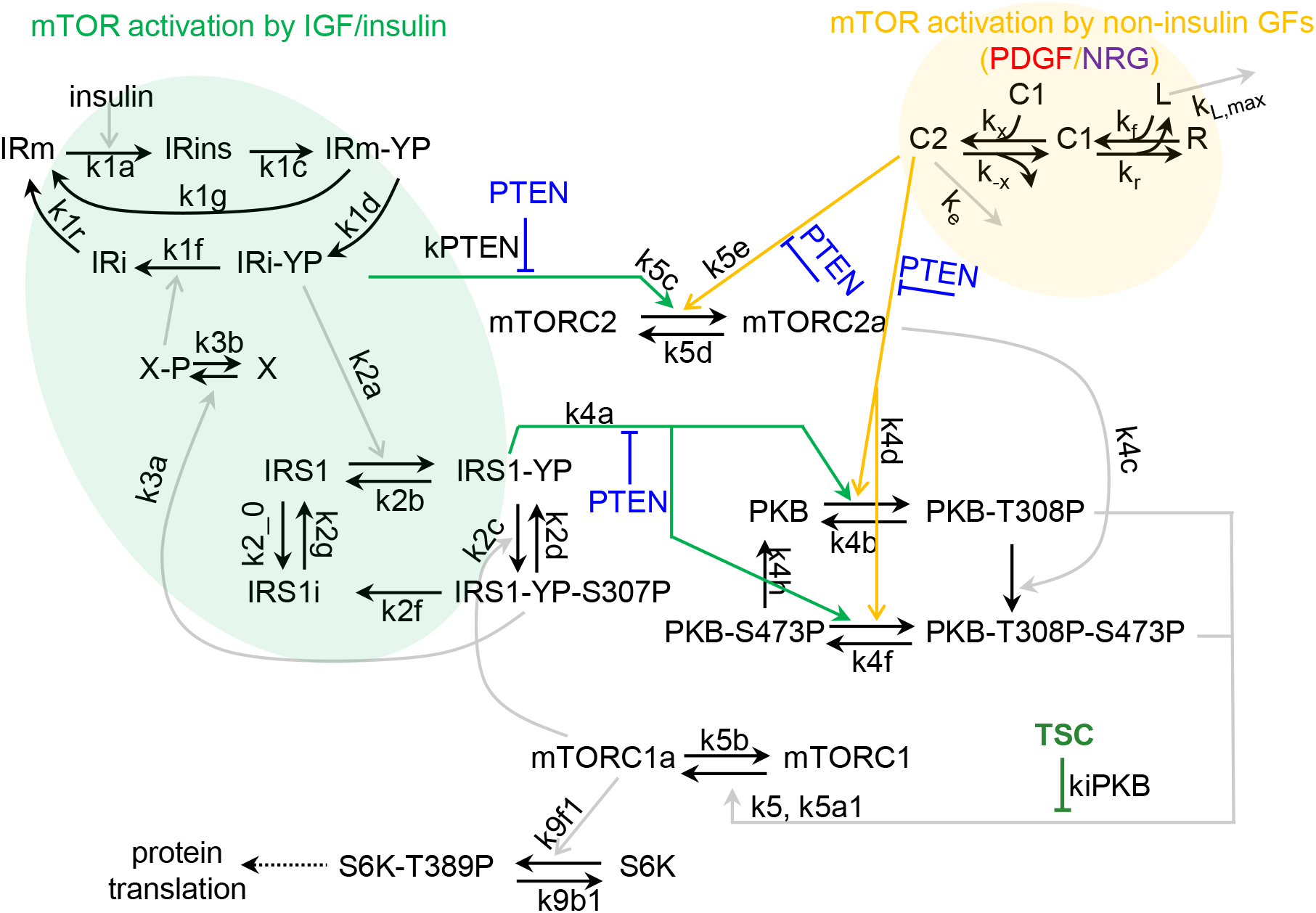
The reaction scheme of the reference model of canonical mTOR pathway. The model is based on integration of (7, 8) models of insulin-dependent and independent signalling. mTOR activation by insulin-like ligands starts from phosphorylation of IR and IRS1. IRS1 negatively regulates IR through a feedback mediated by hypothetical phosphatase X, dephosphorylating IRi-YP (7, 11). Activated mTORC1 (mTORC1a) negatively regulates IRS1 activity through Ser phosphorylation. mTOR activation by non-insulin ligands (L; PDGF or NRG) starts from the formation of ligand (L)-receptor (R) monomeric C1 complex, followed by the formation of active dimeric C2 complex. mTORC2 is activated by C2 and Tyr-phosphorylated IR (IRi-YP). PKB is phosphorylated at Tyr308 by C2 and Tyr-phosphorylated IRS1 (IRS1-YP), and at Ser473 by activated mTORC2. Tyr-phosphorylated PKB activates mTORC1, following by S6K phosphorylation. The transitions between model variables are shown by black solid lines, and enzymatic reactions are shown by coloured lines.

### 2.2. Four components are predicted to be potent regulators of mTOR pathway activity

We next explored the effects of changes in the abundance of mTOR components on overall activity of mTOR pathway through randomised variations in abundance of mTOR components in the reference model. Changes in the abundance of individual variables differentially affected the model output, with linear changes in S6K activity upon 10-fold increase/decrease in abundance of TSC, S6K, et. cet. (Figs. S4A). To determine the most influential variables with highest combined effect of the system, we analysed the correlation between S6K activity and functions combining abundances of several variable (determined as the multiplication of abundances of the activating components, dividing by abundances of the inhibiting components PTEN and TSC). We found that the combined abundance of 4 components, PKB, S6K, TSC, PTEN had the highest correlation with S6K activity (R=0.94 for non-insulin and R=0.87 for insulin stimulus, Fig. 2A), while combinations of less correlated variables had lower effect on S6K activity (Figs. S4,S5). Correlation of S6K activity with other pathway components (receptors, IRS1) was low (Fig. S4A) and their addition to combined PKB, S6K, TSC and PTEN abundances did not increase its correlation with S6K activity (Fig. S4B). This suggests that modulation of the abundance of these 4 components would be an efficient way to regulate the mTOR pathway activity.

**Figure 2.**
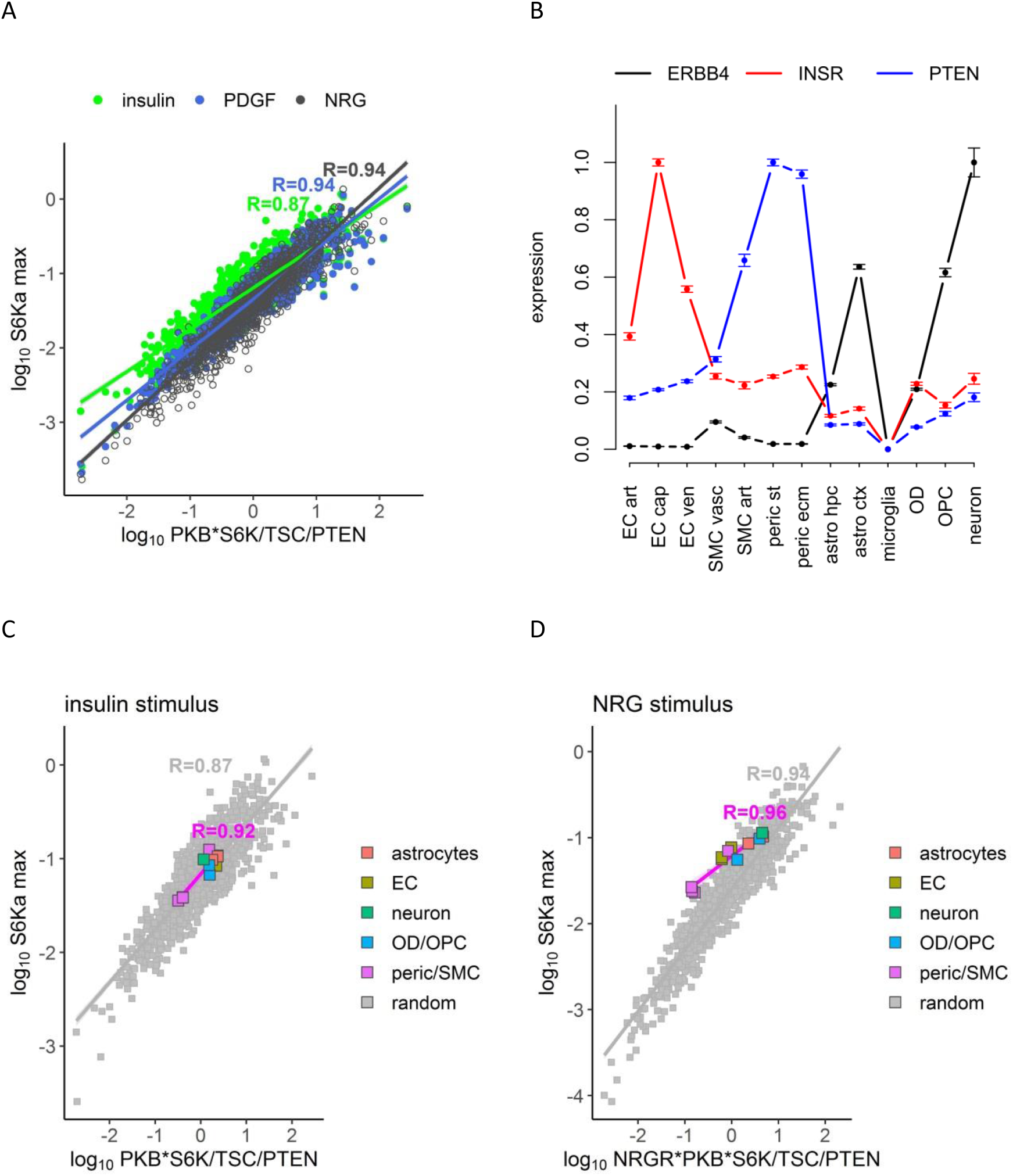
Variations in abundances of mTOR pathway components affect mTOR pathway activity. **A**. Linear correlation between S6Ka and a combination of PKB, S6K, TSC and PTEN abundances in 1000 simulated cells with random variations in abundances of mTOR components. Colours indicate different stimulus. **B**. Example expression of mTOR pathway components in human brain. The data are based on snRNA-seq of cerebrovascular human tissue (6). ERBB4 and INSR are examples of NRG and insulin receptors. Expression is averaged between brain samples from 8 healthy donors and normalized to maximum. Error bars show SEM. Designations are: EC - endothelial cells (arterial, capillary and venous); SMC - smooth muscle cells (vascular, arterial); peric - pericytes (solute transport and ECM-regulating); astro -astrocytes (hippocampal and cortex); OPC - oligodendrocyte precursor cells and OD - oligodendrocytes. **C,D**. Comparison of changes in S6Ka upon random (grey) and brain cell-specific (coloured) variations in the abundance. Correlation plots (A,C,D) show linear correlation between log_10_ of S6Ka and indicated functions of abundances, highly correlating with S6Ka. The maximum of S6Ka was calculated after 2h of stimulation with 10 nM of the indicated ligand. Different subtypes of cells (astrocytes, EC, mural cells (pericytes, SMC) and OD/OPC) are grouped for simplicity.

### 2.3. PTEN expression is the key determinant of brain cell-type specific variation in mTOR pathway activity in response to insulin

Analysis of expression of the mTOR pathway components in the previously identified cell types revealed large variability in expression of ligands and their receptors. Hence insulin and IGF1 receptors (INSR, IGF1R), PDGF and NRG receptors (PDGFRA/B, ERBB4) (Fig. 2B, Fig. S3; (6)) expression levels were > 10-fold different between cell types. The mTOR pathway intracellular regulatory components on the other hand, such as PKB, TSC and S6K showed more consistent expression between cell types, except for PTEN, which also showed wide variance in expression (Fig. S6). Thus, PTEN represents the intersection between the most influential model variables and biological variance within the brain.

We next used our model with abundances of mTOR components derived from their normalized average gene expression in each type of brain cells (Supplementary Information; Table S2; (12)). With respect to the insulin signalling pathway, similar to the model with randomised abundances, the combined abundance of the four components - PKB, S6K, TSC and PTEN had highest correlation with S6K activity in brain cells (Fig. 2C), while variations in IR and IRS1 levels had less effect (Fig. S5). Amongst the four components PTEN abundance had the largest variations and, therefore, is likely the main determinant of S6K activity, in particular in the mural cells where PTEN is the most divergent from the average gene expression (Fig. S6).

PDGF and NRG stimulation were both strongly affected by their receptor abundance (Fig. 2D, Fig. S5), which showed more drastic variations between the cell types compared to other mTOR components (Fig. S6). Thus, the cell-specific activation of mTOR by non-insulin ligands is regulated by receptor abundance and, to a lesser extent by PKB, S6K, TSC and PTEN abundance.

### 2.4. mTOR activation is brain cell type and ligand specific

We next analysed mTOR activation in a broad range of physiologically relevant (0.01-1 nM) ligand concentrations (13, 14) to assess the relative sensitivity of different cell types to different ligands. The model predicts > 10-fold differences in mTOR activities between cell types and ligand-specific mTOR activation (Fig. 3A,B). Thus mTOR in neurons, oligodendrocyte precursor cells and vascular smooth muscle cells were responsive to all stimuli; mTOR in pericytes and arterial smooth muscle cells were mainly responsive to PDGF; mTOR in astrocytes, endothelial cells and oligodendrocytes were responsive to NRG and insulin; and microglia had low activity of mTOR pathway upon all stimuli. The differential responsiveness was principally driven by the abundance of non-insulin receptors and 4 key variables (PKB, S6K, TSC and PTEN), as described above.

**Figure 3.**
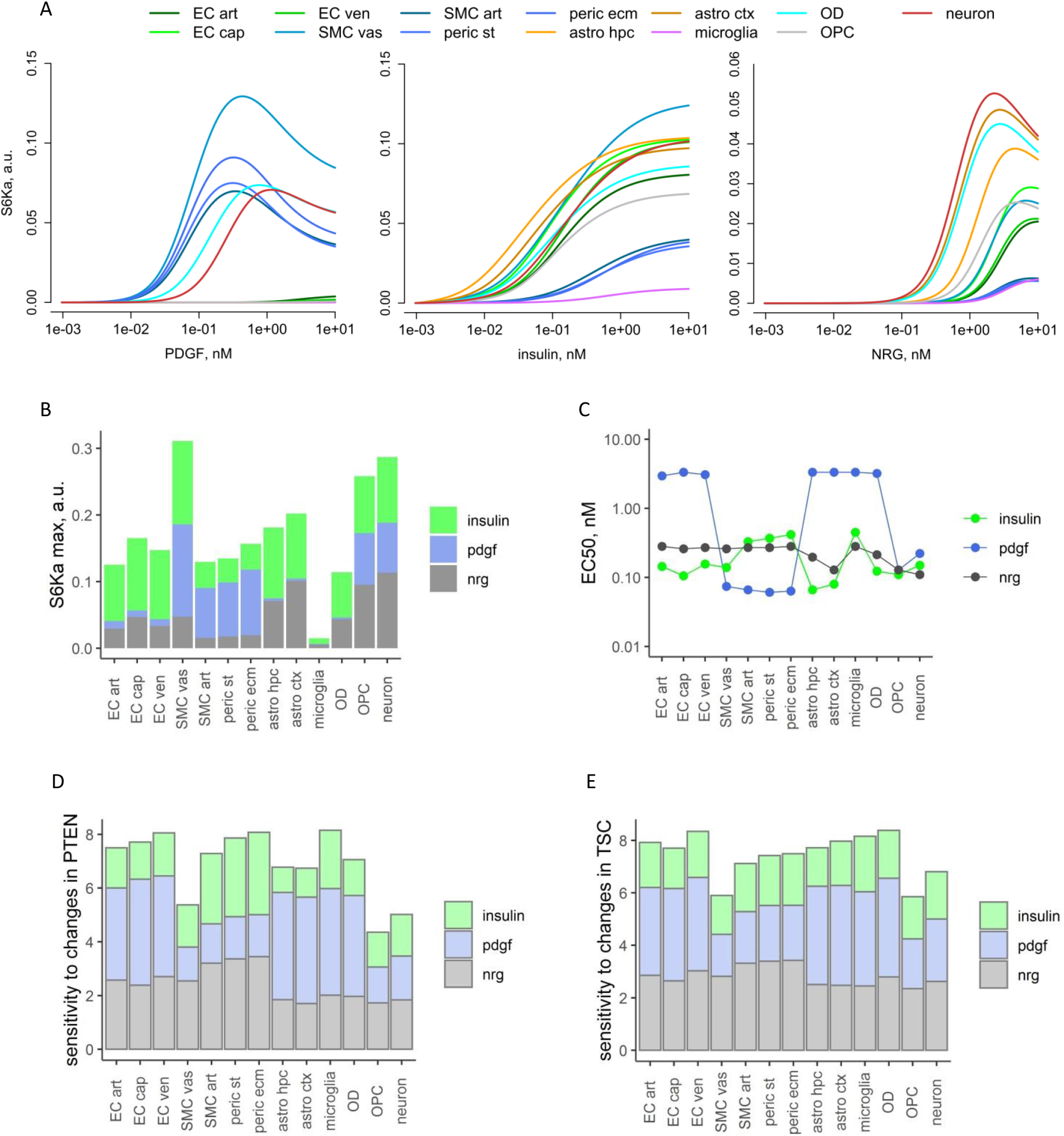
Modelling of mTOR pathway activity in different types of brain cells. **A**. S6Ka levels after 2h of mTOR stimulation with different doses of ligands (after 10 h without ligands). The SK6a levels (with background S6Ka being subtracted) in different cells are indicated by colours. **B,C**. Cell-specific changes in maximal S6Ka (B) and EC50 (C) in the range of ligand concentrations ≤ 10 nM. For NRG stimulation S6Ka was multiplied to 2 and EC50 divided to 5 for easy comparison. **D,E**. The sensitivity of mTOR pathway to variations in the abundance of PTEN (D) or TSC (E). The sensitivity was defined as the fold change in maximal S6Ka upon 4-fold changes in PTEN or TSC abundance (2-fold decrease vs 2-fold increase).

We also calculated the effective dose of ligands required for 50% stimulation (EC50) and found this varied substantially between cell types (Fig. 3C). We found that cells exhibiting a low EC50 to a ligand, typically had high maximal S6K activity (“high mTOR cells”) and cell types with high EC50 had lower maximal S6K activity (“low mTOR cells”). Hence high mTOR cells, such as pericytes upon PDGF stimulation or neurons upon NRG stimulation, are expected to show a large increase in S6K activity even with low ligand concentration. In contrast, “low mTOR” cells, for example endothelial cells upon PDGF stimulation or pericytes upon insulin stimulation have low maximal S6K activity and, typically, high EC50, and therefore likely to remain inactive even with higher ligand concentrations (Fig. 3B,C). Therefore, variations in S6K activity and EC50 between the cells provide a mechanism, whereby over a range of ligand concentrations, the mTOR responses can remain cell type specific.

### 2.5. Sensitivity of the mTOR pathway to changes in PTEN or TSC

PTEN and TSC are two important negative regulators of the mTOR pathway and their abundances in brain cells have been found to be altered under several pathological conditions or secondary to inherited mutations. Thus, PTEN is reduced in many types of cancer, while TSC1 or TSC2 mutations cause Tuberous Sclerosis through presumed loss of function (15, 16). To explore cell-specific effects of variations in the levels of these key mTOR regulators, we calculated fold changes in S6K activity upon 4-fold changes in the level of PTEN or TSC (Fig. 3D,E).

We found that insulin-dependent mTOR activation was resilient to alterations in PTEN or TSC due to the presence of negative feedback loops in the insulin-related part of the model, stabilizing the mTOR pathway upon perturbations (5). Nevertheless, mTOR responses to insulin-like ligands in pericytes and arterial smooth muscle cells were affected by changes in PTEN to higher extent (3-fold difference in S6Ka) than in TSC (< 2-fold), due to high levels of PTEN in these cells. With respect to PDGF signalling, both PTEN and TSC abundance had pronounced effects (4-fold) on endothelial cells, astrocytes, microglia and oligodendrocytes in our model. However, the levels of PDGF receptors were negligible in microglia and astrocytes, leaving endothelial cells and oligodendrocytes as the most vulnerable to TSC and PTEN heterogeneity and spontaneous mutations. The changes in PTEN and TSC had less effect (2-fold) on smooth muscle cells, pericytes and neurons upon PDGF stimulation. Finally, in NRG-mediated mTOR activation, changes in TSC affected all cell types (~3-fold), whilst changes in PTEN mainly affected mural cell types (3.5-fold).

### 2.6. Alzheimer’s Disease selectively affects insulin-mediated mTOR pathway activity

We next investigated how Alzheimer’s Disease (AD) might affect cell type specific activity of mTOR pathway. The comparison on the abundance of mTOR components under healthy and AD conditions demonstrated relatively minor changes in multiple mTOR components (Fig. S6). Therefore, we used the model to explore the potential combined effects of these changes on overall activity of mTOR pathway.

The maximal activity of S6K in response to insulin was lower in AD conditions for endothelial cells, vascular smooth muscle cells and hippocampal astrocytes; and in response to PDGF the S6K activity was reduced for vascular smooth muscle cells, pericytes and neurons (Fig. 4A). Interestingly, activation of S6K by insulin was stimulated in two types of AD cells - microglia and oligodendrocytes, mostly due to higher expression of PKB and S6K (Fig. S6).

**Figure 4.**
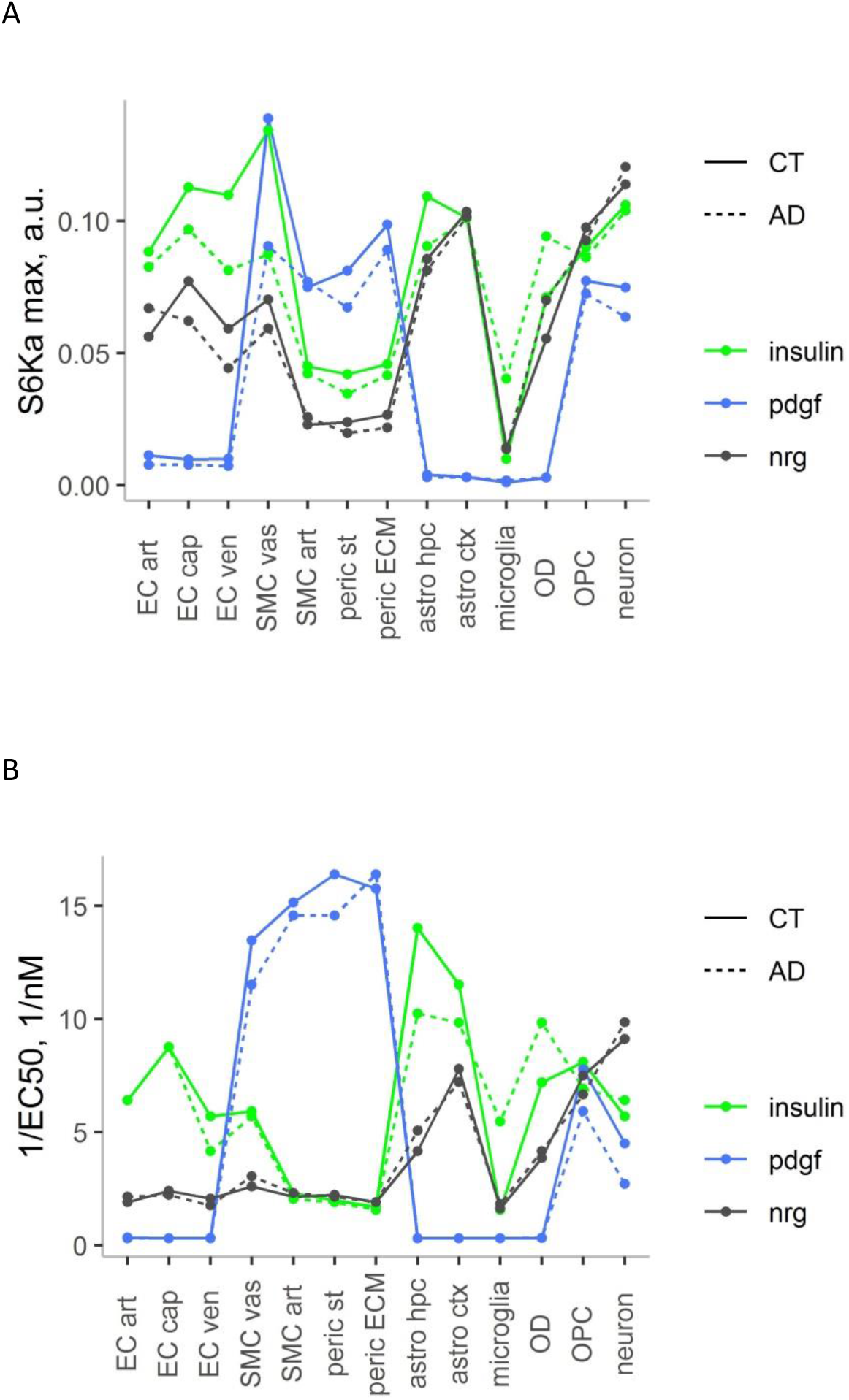
Comparison of mTOR activation in brain cells between healthy and AD conditions. Maximal S6K activity (**A**) and the potency of the ligands (reciprocal of EC50) (**B**) were calculated for mTOR activation in healthy (control, CT) and AD conditions, as shown in legends. The calculations were done using averaged abundance of mTOR components in brain samples from 8 healthy or 9 AD donors (6). The maximal S6Ka was calculated after 2h of mTOR stimulation with various concentrations of ligands ≤ 10 nM. The background activity was subtracted. The responses to insulin, PDGF and NRG are shown by different colours.

The reciprocal of EC50 reflects the potency of the ligands towards mTOR activation (Fig. 4B). The potency of NRG was largely unchanged in AD, and PDGF potency showed a minor decrease in smooth muscle cells, solute transport pericytes and neurons. However, the potency of insulin was noticeably decreased in astrocytes and increased in microglia and oligodendrocytes. The simultaneous increase of maximal S6Ka and insulin potency in microglia and oligodendrocytes means that these cell types are more responsive and have a greater capacity to increase mTOR pathway activity. Microglia are known to be functionally activated during the AD (17), which agrees with the predicted upregulation of mTOR in microglia, whilst the change in oligodendrocytes is novel.

## 3. Discussion

Whilst the mTOR pathway is a ubiquitous regulator of cell metabolism and growth, potential heterogeneity in expression and activity in brain cells has not been well explored. Characterisation of heterogeneity is important since there are a large number of disorders, including neurological disorders, where a central feature is alteration in mTOR pathway activity, but the cell types involved in disease manifestation are highly variable. This includes genetic disorders such as Tuberous Sclerosis arising from loss of function in TSC1 or TSC2 (16), as well as late onset degenerative conditions such as Alzheimer’s Disease, where multiple lines of evidence implicate an important role for mTOR in disease progression (18, 19). Furthermore, in many cancers somatic mutations in the mTOR pathway genes contribute to tumorigenesis (20).

Perhaps unsurprisingly investigation of changes in mTOR in these disorders can show contradictory results. This is likely due to the complexity of the mTOR pathway, with multiple upstream signalling mechanisms, as well as positive and negative feedback loops. Such a system may display non-linear behaviours and in order to identify any emergent properties it is necessary to model key aspects of the pathway. Previously published mathematical models of the mTOR pathway have been limited to include mTOR activation in isolated cell lines. We have now established a more comprehensive model of mTOR activation in the whole brain, incorporating three major signalling pathways, operating in various types of brain cells. Our model describes the heterogeneity of mTOR expression in brain cells, based on a large publicly available snRNA-seq dataset (6). Using such large datasets with hundreds of thousands of nuclei/cells allows reproducible measurements of relatively low inter-donor variations in gene expression between different cell types (Fig. S6).

The model represents the first combination of mathematical modelling with snRNA-seq data to model mTOR dynamics in different types of brain cells, and has several limitations. Thus, we only describe the canonical mTOR pathway, operating in mature, differentiated brain tissues, where nutrients are non-limiting. We included three groups of receptors (IR, PDGFR and NRGR) that are highly expressed in the brain. The model can be further extended by including other groups of receptors and processes, e.g., the MAPK pathway, which modulates mTOR in differentiating cells. We have simplified the representation of each group of ligands as a single entity, thus modelling the cumulative effects of ligand groups on brain cells. Future iterations could refine the model by separately considering the individual ligands in each group. Another limitation of our model is that the abundance of mTOR pathway components was estimated from transcriptional expression data. Whilst, mRNA levels are key determinants for protein abundances, translational regulation and often unknown post-translational modification will also alter protein activities (12). And finally, the model employs independent descriptions for mTOR activation by each ligand, which can be further extended by adding the competition between ligands for downstream mTOR components, to describe simultaneous activation of mTOR by different ligands.

Despite these limitations we believe our model provides new insights on cell-type specific mTOR activation by different upstream activators. We found large variations in responsiveness (effective dose) and magnitude of mTOR activation between cell types, which was also dependent on the stimulating ligand. Vascular smooth muscle cells, oligodendrocyte precursor cells and neurons were predicted to be readily activated by all the ligands (Fig. 5) due to relatively high abundance of most mTOR components and reduced levels of PTEN or TSC. In contrast, endothelial cells were specifically sensitive to insulin and pericytes to PDGF, which is consistent with their specific location and, thus, exposure to those stimuli. Endothelial cells are the key interface between blood and brain, and insulin synthesized and released into the systemic circulation from the pancreas and is well placed to regulate metabolism of endothelial cells in the brain. Pericytes, on the other-hand, are on the abluminal side of the blood-brain-barrier adjacent to endothelial cells. They are more likely to be exposed to higher levels of PDGF secreted by endothelial cells (21) or neurons (22), in order to respond signals from the vascular or brain compartment, respectively. We found, that different subgroups of the same cell types, such as sub-populations of endothelial cell, pericytes or astrocytes have similar characteristics of the mTOR pathway activity (Fig. 5). The only exception was vascular smooth muscle cells, showing intermediate characteristics between EC and smooth muscle cells/pericytes, which reflects the differences in the abundance of mTOR components between these cell types (Fig. S6).

**Fig. 5.**
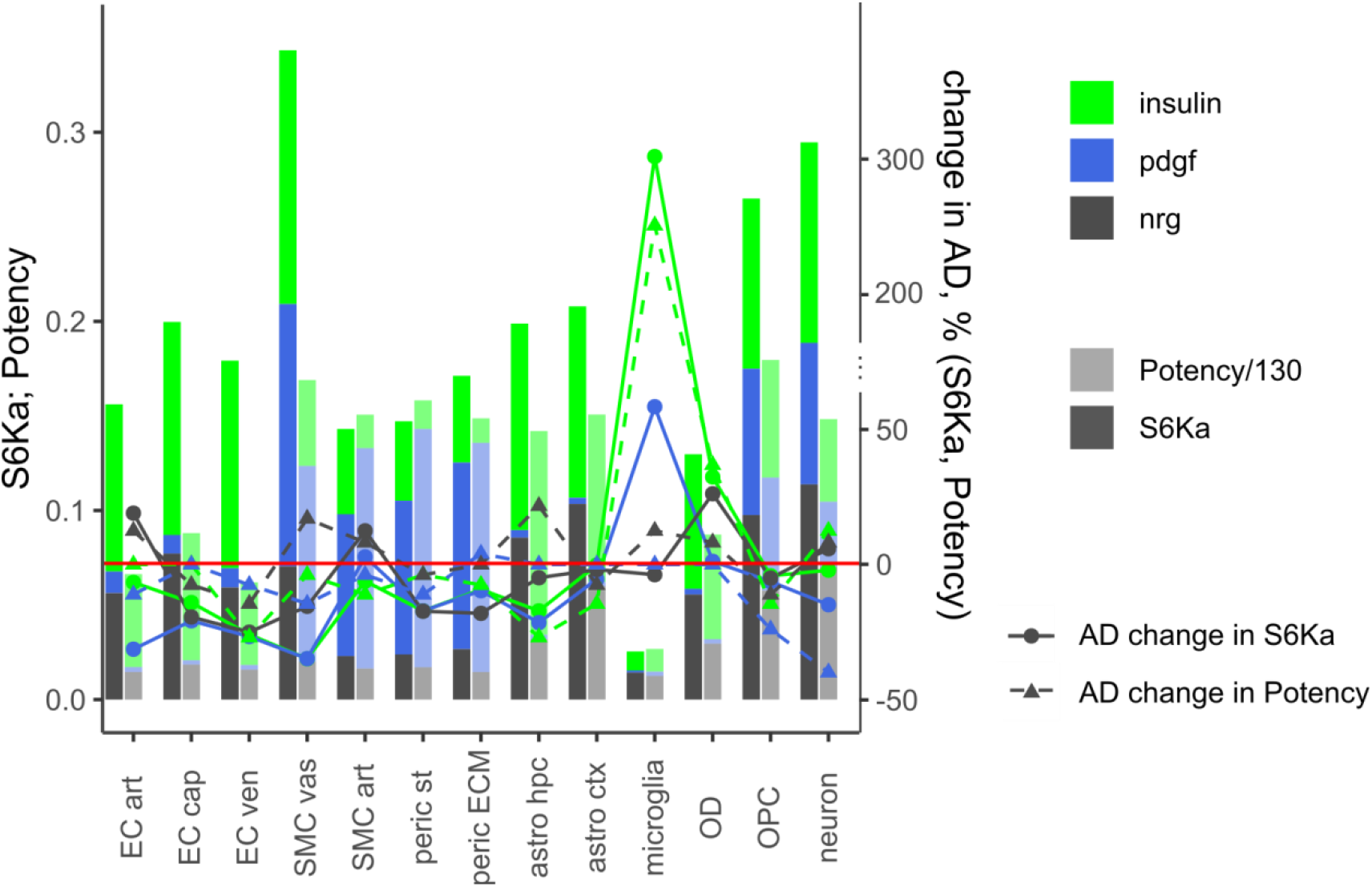
Summary of the differences in mTOR activities between healthy and AD cells. The bars show maximum of S6Ka (brighter colours) and the potency (1/EC50/130, lighter colours) in different types of the brain cells. Stimulation with different ligands is indicated by colours. The lines show % changes in the maximal S6Ka and potency in AD vs control. Changes in S6Ka and potency are shown by circles (connected by solid lines) and triangles (dashed lines), respectively. Red line indicates no changes. The changes are calculated as the difference between AD and control values divided by the control values.

Our analysis indicated that insulin-related activity of mTOR pathway is not strongly regulated by the abundance of the insulin receptor. This may be due to the presence of negative feedbacks on insulin receptor activity, overriding potential effects of changes in IR and IRS abundance. Instead, we found that the main determinants of variation in mTOR pathway activity in response to insulin are PKB, the negative regulators - PTEN and TSC, and the final effector S6K. This is consistent with findings from previous models, which proposed PKB and PTEN as key regulators of mTOR pathway activity (23, 24). TSC is a well-known central regulator of mTOR pathway, mediating multiple signalling inputs into the mTOR pathway (25). Surprisingly then, when we next modelled the impact of reduction in PTEN and TSC, we found that insulin stimulated mTOR was in general resilient to such reductions. PDGF and NRG stimulated mTOR, on the other hand, showed strong dependence on their receptor abundance, as well as PTEN and TSC. However, different cell types showed sensitivity to reductions in PTEN or TSC, dependent on whether it was PDGF-stimulated mTOR (endothelial cells, astrocytes, microglia, oligodendrocytes) or NRG-stimulated mTOR (smooth muscle cells and pericytes).

Hence, our model highlights the complex interaction between ligand, mTOR negative regulators and the cell type. Our findings may provide insight into why particular cell types are preferentially affected by particular diseases. We speculate that where a combination of genetic mutation and the dominant signalling molecules leads to enhanced activity of mTOR pathway, this leads to increased protein biosynthesis in particular cell types. For example, in Tuberous Sclerosis, an overgrowth or abnormalities of astrocytes are pathognomonic of the condition. Furthermore, our modelling identifies PDGF signalling as a driver of mTOR when TSC is deficient, which suggests that blocking PDGF signalling could be therapeutically beneficial. In cancer, somatic mutations may lead to loss of PTEN or TSC function (20) and, again, PDGF may be an important driver of cancer growth. In this case PDGF receptor monoclonal antibodies have been proposed as a promising cancer therapy (26).

Using the published snRNA-seq data we were also able to investigate the impact of AD on cell-type specific activity of mTOR pathway. AD did not significantly alter responses to NRG stimulation, and responses to PDGF were only affected (reduced) in smooth muscle cells and pericytes. However, the insulin signalling pathway was selectively altered, which is consistent with the concept of AD as ‘Type III Diabetes’ (27). Our model identified increased potency of insulin signalling in microglia and oligodendrocytes, coinciding with an increased maximal S6K activity in these cells. The predicted upregulation of mTOR efficiency in oligodendrocytes would counteract the myelin damage, which is typically observed during the AD (17). And the predicted upregulation of mTOR efficiency in AD microglia is consistent with hyper-activation of inflammatory functions in AD. It is possible that such changes are either part of the disease mechanism to promote disease progression or part of a protective response. Hence, agents which further modulate insulin signalling in AD may represent therapeutic opportunities.

We conclude that the mTOR pathway is differentially regulated in brain cells, affecting brain functions under normal and pathological conditions. The complexity of the pathway necessitates mathematical modelling to fully appreciate the impact of perturbations.

## Data accessibility

The R code of the model is freely available at https://github.com (the full URL is https://github.com/alex297/mTOR; read-me file is provided in the Supplementary Information).

## Authors’ contributions

ZC conceived the study, AP built and analysed the model, all authors contributed to the study design and paper writing

