## Supplementary Information for "Mathematical modelling of brain mTOR activity identifies selective vulnerability of cell types and signalling pathways"

**List of content**


1. Reference model p.1
   1.1. Model description p.1
   1.2. Incorporation of the negative feedback from mTOR to IRS1 p.2
   1.3. Model assumptions p.3
   1.4. ODE equations and data fitting p.3
   1.5. Selection of GFs to be included in the model p.8
   1.6. Simulations of random fluctuations in the abundances of mTOR components p.10
2. Modelling the mTOR pathway in brain cells p. 13
3. References p. 19


### 1. Reference model

#### *1.1. Model description*

The model describes the kinetics of canonical AKT/mTOR pathway, starting from the activation of tyrosine kinase receptors by their GFs. The input of our model is a stepwise-added extracellular GF concentration and the main output is the S6K activity (S6Ka). To describe mTOR activation by both insulin-like and non-insulin types of GFs, we combined published models of insulin and PDGF signalling (1, 2). These models were fitted to kinetic data at various steps of the mTOR pathway activation pathway, such as phosphorylation kinetics of receptors, IRS1 and PKB (AKT). The insulin pathway represents the basis of our model and includes Tyr phosphorylation and activation of insulin receptor (IR) after its binding to insulin/IGF (formation of membrane-bound IRm-YP); internalization of IRm-YP, resulting in the formation of active IRi-YP; inactivation/dephosphorylation of IRi-YP by hypothetical phosphatase X through IRS1-mediated negative feedback (1, 3); Tyr phosphorylation and activation of IRS1 by IRi-YP (formation of IRS1-YP; Ser phosphorylation of IRS1-YP by activated mTORC1 (mTORC1a), resulting in its inactivation (mTOR-mediated negative feedback, described below); Tyr phosphorylation and activation of PKB by IRS1-YP; activation of mTORC2 by IRi-YP; Ser phosphorylation of PKB by activated mTORC2 (mTORC2a); activation of mTORC1 by Tyr-phosphorylated forms of PKB, and, finally, phosphorylation of S6K by mTORC1a ((1), Fig. 2). For non-insulin GFs the initial steps include the formation of monomeric (C1) and dimeric (C2) complexes between PDGF (or NRG) ligands (L) and their receptors (R) and phosphorylation of PKB and mTORC2 by C2 (Fig. 2).

The model was extended by including the inhibitors PTEN and TSC, which play a key role in regulation of the mTOR pathway under various pathological conditions (4, 5). PTEN inhibits steps of the pathway upstream of PKB activation (described in details below), while TSC inhibits the final steps of mTOR activation downstream of PKB (Fig. 2). The mechanisms of the background activation of mTOR observed in absence of GFs (1) are largely unknown. In our model, we simply included low background phosphorylation of PKB, replacing IR/IRS-dependent background activation of mTOR in the insulin model. All simulations were done after 10 h of background stimulation of mTOR without GFs, starting from zero initial conditions for all phosphorylated species. Model parameters were predominantly taken from published models (Table S1). Remaining parameters (including parameters combining the models together and related to the negative feedback) were manually adjusted to the published data. The data used for fitting included time courses of activation of different variables with varying concentrations of insulin (Fig. S2A), PDGF (Fig. S2B) and NRG (Fig. S2C) (1, 2, 6), as well as changes in the activities of mTOR and PKB upon rapamycin and TSC knock out (TSC-KO) (Fig. S1). The description of the model equations and parameters is presented below.

#### *1.2 Incorporation of the negative feedback from mTOR to IRS1*

TSC mutations cause activation of mTOR, while rapamycin treatment inhibits mTOR. These two opposite interventions, covering the wide range of mTORC1 activities allowed us to explore the feedback structure of the mTOR pathway (7-9). Available data suggests that, similarly to many other cell types (7-9), brain cells respond to TSC-KO by downregulating PKB activity (by reducing phosphorylation levels) and respond to rapamycin by increasing PKB activity, suggesting the presence of a negative feedback loop (10). The feedback is mediated by Ser phosphorylation of IRS1 by mTORC1a, resulting in IRS1 and IR inactivation (7). We replaced the positive feedback in the insulin model (1), which was suggested to operate mainly in adipocytes (11), with the negative feedback, which is supported by multiple sources (7-10). The negative feedback was implemented by minor changes to the insulin model. We omitted the reactions of PKB phosphorylation by IRS1-YP-Ser and replaced IRS1-YP with IRS1-YP-Ser in the activation of X phosphatase, in agreement with experimental observations on the negative regulation of IRS1 and IR by mTORC1a (7). We also added the basal phosphorylation of X with the rate constant k3_0 to restrict IR activation upon rapamycin treatment (in absence of IRS1-YP-Ser). The inactivated IRS1-S307P was renamed into IRS1i (Fig. 2). The reaction of Tyr phosphorylation of PKB-Ser by IRS1-YP was retained from the full reaction scheme from (1), with the respective rate constant k4a fitted to PKB timecourses. These changes allowed us to simultaneously describe the kinetics of mTOR components (Fig. S2) and the response to rapamycin and TSC-KO (Fig. S1). Rapamycin was modelled by setting the rate of mTORC1 activation by PKB to zero. The model describes the observed ~ 2-fold reduction of PKB-Ser levels in TSC-KO upon insulin treatment, as well as ~ 2-fold increase in PKB activity upon rapamycin treatment (Fig. S1, (8, 9)). There was also an increase in background mTORC1 activity in absence of insulin in TSC-KO compared to the wild type (WT), in agreement with the data (8, 9). The mTORC2a was not significantly affected in TSC-KO, in agreement with published observations (8). ODEs were solved using deSolve package in R (12).

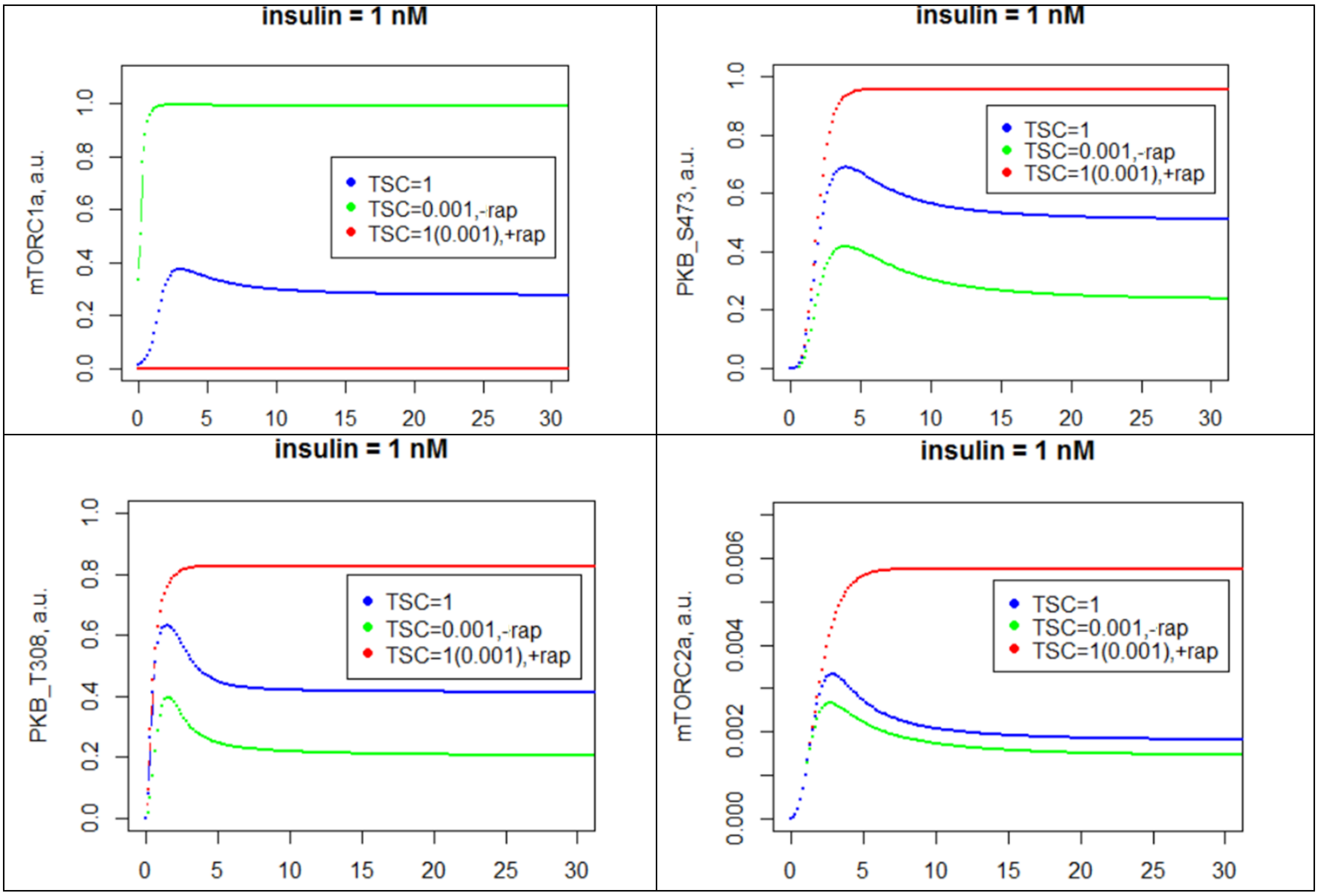


Figure S1. Effects of mTORC1a-mediated negative feedback on the timecourse of mTOR stimulation by 1 nM insulin. Blue lines correspond to the WT, green lines - to TSC-KO, which was modelled by 1000-fold reduction in TSC level (TSC=0.001). Red lines correspond to rapamycin (rap) treatment (not affected by the TSC levels).

#### *1.3. Model assumptions*

1) mTOR activation is involved in multiple processes: growth, differentiation, stress response, etc., with multiple pathways participating in mTOR activation/inhibition (13). In our model we consider only the canonical mTOR pathway, operating in mature, differentiated brain cells under normal, unstressed conditions with sufficient nutrients.
2) The reference model integrates previous models of the mTOR pathway, which were developed for non-brain cell lines (adipocytes, fibroblasts). We assumed that the main structure of the pathway and kinetic parameters are preserved between different tissues. However we modified the previous models to ensure mTOR components (inhibitors, GFs) and processes (negative feedback loops) relevant to the brain were included in the model.
3) In the brain-specific model the levels of mTOR components were equal to normalized sum of expression of respective genes taken from our snRNA-seq data. Therefore, the model describes the cumulative, averaged effect of each group of GFs.
4) In brain-specific model we assumed that mRNA levels of mTOR pathway components correlate with their protein levels (14). Since the protein levels are largely unknown, the model predicts not the absolute, but relative differences in mTOR pathway activity between the cells. The levels of mTOR components in the adapted published models are also expressed in relative units, therefore the model parameters (e.g., reaction rates) are relative.
5) The activation of mTOR pathway by each of the three brain-specific groups of GFs (IGF, PDGF, NRG) was considered separately from each other, therefore we ignored the competition between different ligand-receptor complexes for the downstream components of the pathway.

#### *1.4. ODE equations and data fitting*

Similarly to the adapted models (1, 2), the concentrations of all mTOR components in our model are dimensionless; the time unit is a minute; and the total/max concentration of each component of the reference model is equal to 1. The variables of the insulin model were rescaled from maximal levels of 100% (1) to 1, with respective changes in model parameters. The concentrations of insulin/IGF and other GFs are expressed in nM.

*The insulin-related part of the reference model*

The insulin-related part of the model was fitted to the temporal kinetics of mTOR components after stimulation with 10 nM insulin (Fig. S2A; (1)), as well as upon TSC-KO and rapamycin treatment (Fig. S1). The simulated time courses of different variables were very close to previously obtained time courses (1), reflecting similar kinetical properties of our and previous models. Some of the simulated time courses (e.g., for IR, IRS) showed very good fit, while fitting of other variables (e.g., PKB_T308) by both models was less efficient, possibly due to the large noise in the data (fewer data points; Fig. S2A).

Different forms of IR receptor are: IRm is inactive; IRins is IR bound to insulin, non-activated; IRm-YP is Tyr phosphorylated membrane receptor complex; IRi-YP is Tyr phosphorylated and internalized active receptor complex (main active species in (1) model); IRi is non-active dephosphorylated complex. Different forms of IRS1 are: IRS1 - inactive; IRS1_YP - Tyr phosphorylated, active; IRS1_YP_S307P - Ser 307 phosphorylated and active towards phosphorylation of the phosphatase X; and IRS1i - Ser phosphorylated and non-active. The IR-related equations are:

$$\frac{dIRm}{dt} = k1r \cdot IRi - k1a \cdot IRm \cdot insulin + k1g \cdot IRm\_YP$$

$$\frac{dIRins}{dt} = k1a \cdot IRm \cdot insulin - k1c \cdot IRins$$

$$\frac{dIRm\_YP}{dt} = k1c \cdot IRins - k1d \cdot IRm_{YP} - k1g \cdot IRm\_YP$$

$$\frac{dIRi\_YP}{dt} = k1d \cdot IRm\_YP - k1f \cdot IRi\_YP \cdot X\_P$$

$$\frac{dIRi}{dt} = k1f \cdot IRi\_YP \cdot X\_P - k1r \cdot IRi$$

Similarly to (1), each measured specie x is scaled by a scaling parameter ky_x:

$$measured\_IR\_YP = ky\_IR \cdot (IRm\_YP + IRi\_YP)$$

*ky_IR=525*

IRS1-related equations are:

$$\frac{dIRS1}{dt} = k2b \cdot IRS1\_YP + k2g \cdot IRS1i - k2\_0 \cdot IRS1 - k2a \cdot IRS1 \cdot IRi\_YP$$

$$\frac{dIRS1\_YP}{dt} = k2a \cdot IRS1 \cdot IRi\_YP + k2d \cdot IRS1\_YP\_S307P - k2b \cdot IRS1\_YP - k2c \cdot IRS1\_YP \cdot mTORC1a$$

$$\frac{dIRS1\_YP\_S307P}{dt} = k2c \cdot IRS1\_YP \cdot mTORC1a - k2d \cdot IRS1\_YP\_S307P - k2f \cdot IRS1\_YP\_S307P$$

$$\frac{dIRS1i}{dt} = k2\_0 \cdot IRS1 + k2f \cdot IRS1\_YP\_S307P - k2g \cdot IRS1i$$

$$measured\_IRS1\_YP = ky\_IRS1\_YP \cdot (IRS1\_YP + IRS1\_YP\_S307P)$$

$$measured\_IRS1\_S307P = ky\_IRS1\_S307 \cdot (IRS1\_S307P + IRS1\_YP\_S307P)$$

*ky_IRS1_YP=58; ky_IRS1_S307=6.7*

$$\frac{dX}{dt} = k3b \cdot X\_P - k3a \cdot X \cdot (k3basal + IRS1\_YP\_S307P)$$

$$\frac{dX\_P}{dt} = k3a \cdot X \cdot (k3basal + IRS1\_YP\_S307P) - k3b \cdot X\_P$$

Different forms of PKB are: PKB - non-active; PKB_T308P - activated by IRS1-YP; PKB_T308P_S473_P - activated by mTORC2a, PKB_S473 - non-active towards mTORC1 activation, (1):

$$\frac{dPKB}{dt} = k4b \cdot PKB\_T308P + k4h \cdot PKB\_S473\_P - \frac{k4a \cdot PKB \cdot IRS1\_YP + k4d \cdot PKB \cdot PI3K}{PTEN + kiPTEN} - k4\_0 \cdot PKB \cdot mTORC2a$$

$$\frac{dPKB\_T308P}{dt} = \frac{k4a \cdot PKB \cdot IRS1\_YP + k4d \cdot PKB \cdot PI3K}{PTEN + kiPTEN} - k4b \cdot PKB\_T308P - k4c \cdot PKB\_T308P \cdot mTORC2a + k4\_0 \cdot PKB - k4c\_0 \cdot PKB\_T308P$$

$$\frac{dPKB\_S473\_P}{dt} = k4f \cdot PKB\_T308P\_S473\_P - k4h \cdot PKB\_S473\_P - k4c\_0 \cdot PKB\_S473\_P - \frac{k4a \cdot PKB\_S473\_P \cdot IRS1\_YP + k4d \cdot PKB\_S473\_P \cdot PI3K}{PTEN + kiPTEN}$$

$$\frac{dPKB\_T308P\_S473\_P}{dt} = k4c \cdot PKB\_T308P \cdot mTORC2a - k4f \cdot PKB\_T308P\_S473\_P + k4c\_0 \cdot PKB\_S473\_P + k4c\_0 \cdot PKB\_T308P + \frac{k4a \cdot PKB\_S473\_P \cdot IRS1\_YP + k4d \cdot PKB\_S473\_P \cdot PI3K}{PTEN + kiPTEN}$$

Here some terms with PI3K function are from the PDGF-related part of the model described below. The inhibition of PKB activation by PTEN is included in the respective steps at the level of PKB activation by IRS1 (for insulin model) or by PI3K (non-insulin model). The inhibition constant *kiPTEN* was estimated from (12), using basal rate of PIP3 degradation, with PTEN scaled to 1. The PTEN inhibition is also included into the activation of mTORC2a (below), based on the existing evidence on the involvement of PI3K in the mTORC2 activation (8, 11). The basal phosphorylation of PKB on Tyr and Ser with the rate constants k4_0, k4c_0 replaced IR/IRS1-dependent basal activation from (1) model. This was done to describe a more general background activation of mTOR observed in experiments with different ligands.

$$measured\_PKB\_T308P = ky\_PKB\_T308P \cdot (PKB\_T308P + PKB\_T308P\_S473P)$$

$$measured\_PKB\_S473P = ky\_PKB\_S473 \cdot (PKB\_S473P + PKB\_T308P\_S473P)$$

*ky_PKB_T308P=4.36; ky_PKB_S473=1.6*

Activation of mTOR complexes was described as:

$$\frac{dmTORC1a}{dt} = \frac{k5}{TSC + kiTSC} \cdot (PKB\_T308P\_S473P + k5a1 \cdot PKB\_T308P) \cdot mTORC1 - k5b \cdot mTORC1a$$

$$\frac{dmTORC1}{dt} = k5b \cdot mTORC1a - \frac{k5}{TSC + kiTSC} \cdot (PKB\_T308P\_S473P + k5a1 \cdot PKB\_T308P) \cdot mTORC1$$

$$\frac{dmTORC2}{dt} = k5d \cdot mTORC2a - \frac{k5c \cdot mTORC2 \cdot IRi\_YP + k5e \cdot mTORC2 \cdot PI3K}{PTEN + kiPTEN}$$

$$\frac{dmTORC2a}{dt} = \frac{k5c \cdot mTORC2 \cdot IRi\_YP + k5e \cdot mTORC2 \cdot PI3K}{PTEN + kiPTEN} - k5d \cdot mTORC2a$$

The mTORC1 activation term includes both forms of Tyr-phosphorylated PKB (*PKB_T308P* and *PKB_T308P_S473P*) and the ratio of PKB activities towards mTORC1 (parameter *k5a1*) was retained from (1). We also included TSC-mediated inhibition of mTORC1a, with total amount of TSC (*TSC)* equal to 1. We assumed that the amount of mTOR activator Pheb is inversely proportional to the amount of non-phosphorylated TSC, which, in turn is proportional to the total amount of TSC and inversely proportional to PKB activity.

The parameter *kiTSC* was chosen to be small enough to describe 2-fold increase in mTORC1a in the TSC-KO relative to WT (Fig. S1). Rapamycin was modelled by multiplying k5 to 0.

The readout of mTORC1 activity is Tyr phosphorylated activated S6K. Equations for S6K are:

$$\frac{dS6K}{dt} = k9b1 \cdot S6K\_T389P - k9f1 \cdot S6K \cdot \frac{mTORC1a}{(km9 + mTORC1a)}$$

$$\frac{dS6K\_T389P}{dt} = k9f1 \cdot S6K \cdot \frac{mTORC1a}{(km9 + mTORC1a)} - k9b1 \cdot S6K\_T389P$$

The rate constants of S6K phosphorylation (k9f1) and basal PKB activation (k4_0, k4c_0) were chosen to give 0.5% activation of S6K in absence of insulin and 12% activation upon 120 min of maximal stimulation with 10 nM insulin (15, 16).

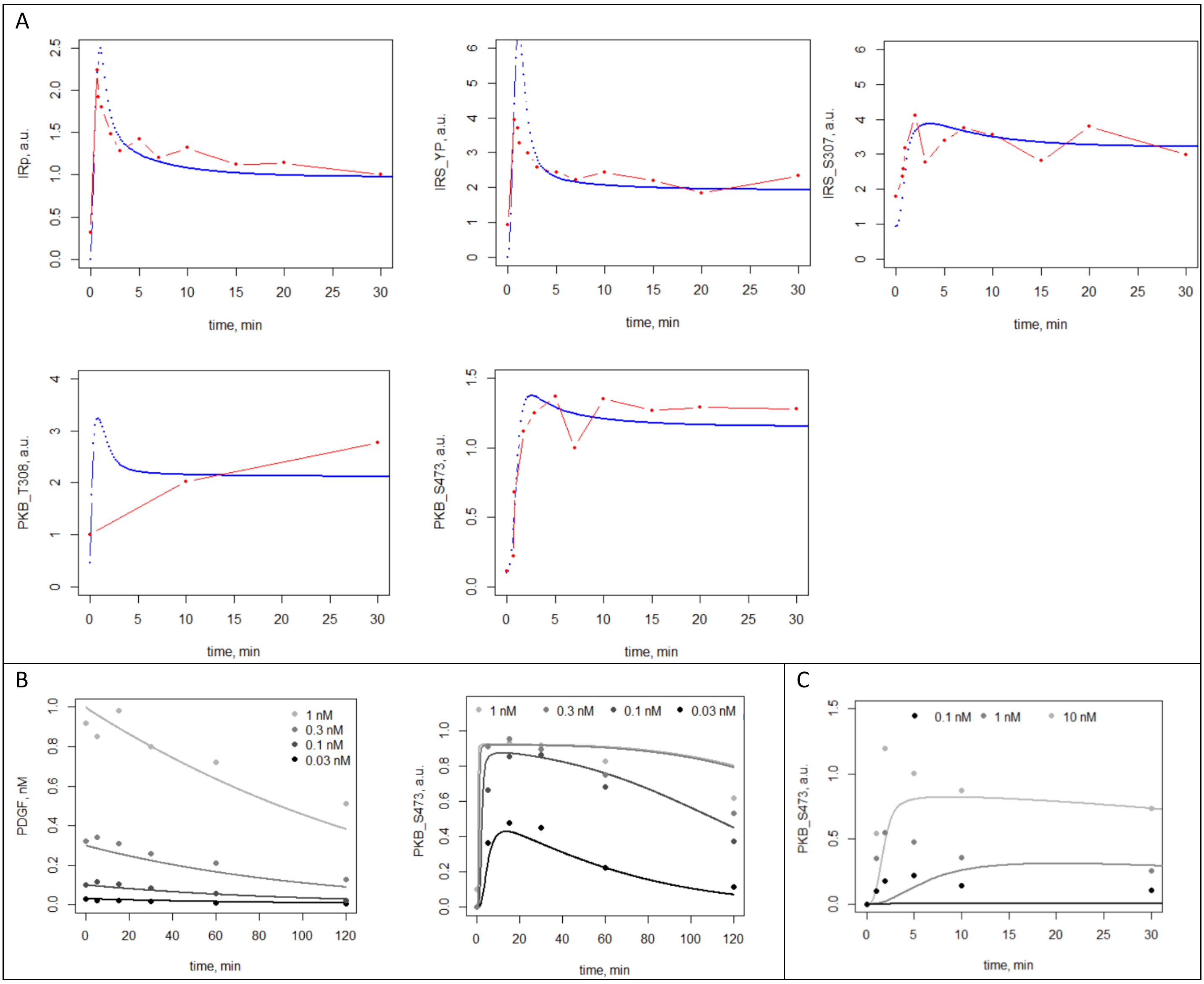

Figure S2. Kinetics of mTOR pathway components upon stimulation with GFs. **A**. Stimulation by 10 nM of insulin. Experimental data points (red) are redrawn from (1). Simulations are shown by blue lines and were done for 10 h without insulin (background stimulation), followed by 30 min with insulin. The data fit corresponds to the original fit (1). **B**. Stimulation with different concentrations of PDGF. Experimental data points are redrawn from (2). Simulations are shown by lines and were done for 10 h without PDGF followed by 2h with PDGF. **C**. Stimulation with different concentrations of NRG. Experimental data points are from (6). Simulations are shown by lines and were done for 10 h without NRG followed by 30 min with NRG.

*Non-insulin (PDGF-related) part of the reference model*

The model of PDGF/Akt signalling was developed in (2). We integrated this model with the insulin model, with some modifications described below. The model describes the formation of ligand-receptor complexes: monomeric ($c_1$) and dimeric complex ($c_2$) (2). The concentration of dimers decreases due to their endocytosis and dissociation from cell surface. The dimensionless concentrations, changing between 0 and 1, for unbound receptor (r), $c_1$ and $c_2$ are described by the equations:

$$\frac{dr}{dt} = k_r c_1 - k_f L r + k_{-x} c_2$$

$$\frac{dc_1}{dt} = k_f L r - k_r c_1 + k_{-x} c_2 - 2 k_x R_0 {c_1}^2$$

$$\frac{dc_2}{dt} = k_x R_0 {c_1}^2 - (k_{-x} + k_e) c_2$$

$$r(0) = 1;\ c_1(0) = 0;\ c_2(0) = 0,$$

where $k_f$ and $k_r$ are the forward and reverse rate constants of the ligand-receptor binding (with the dissociation constant $K_{D,L} = k_r/k_f$); $k_f$ was chosen in the typical range of bimolecular rate constants as 1 $nM^{-1}min^{-1}$ (17), however the choice of $k_f$ does not affect the kinetics, similarly to (2). $R_0$ is the initial dimensionless concentration of receptors; $L_0$ is the initial ligand (PDGF) concentration; $k_e$ is the rate constant of endocytosis; $k_{-x}$ is the rate constant of dissociation of one ligand molecule from $c_2$, releasing one receptor molecule; $k_x$ is the dimerization rate constant.

The kinetics of free ligand is modelled through its sequestration into the receptor complexes, as well as depletion from the extracellular medium (2):

$$\frac{dL}{dt} = R_0 (k_r c_1 - k_f L r + k_{-x} c_2) - \frac{k_{L,max} L}{1 + \frac{L}{K_{M,L}}};\ L(0) = L_0$$

Where $k_{L,max}$ is the depletion rate constant; $K_{M,L}$ is the depletion saturation constant. The initial receptor concentration $R_0$ was fixed in (2), with dimerization rate being proportional to the single parameter $k_x R_0$, while neglecting the depletion of $L$ due to the formation of receptor complexes (assuming that receptor concentration << $L_0$). To model a broad range of potential receptor concentrations, we included the terms describing reactions of association and dissociation of $L$ and $r$, as well of dissociation of $L$ from $c_2$. The parameter $R_0$ was fitted to the (2) data, while preserving the value of $k_x R_0$.

The dimensionless activity of PI3K was previously (2) shown to be fast and described by the following algebraic equation:

$$PI3K(t) = 0.5 \cdot (1 + k_{PI3K} + 2\alpha_{PI3K} \cdot c_2(t) - \sqrt{(1 + k_{PI3K} + 2\alpha_{PI3K} \cdot c_2(t))^2 - 8\alpha_{PI3K} \cdot c_2(t)})$$

Where $\alpha_{PI3K}$ is the estimated receptor/PI3K ratio; $k_{PI3K}$ is the dimensionless receptor-PI3K dissociation constant. To model the dependence of PI3K on changes in receptor levels ($R_0$) between the brain cells, we explicitly included $R_0$ into the parameter $\alpha_{PI3K}$ as $\alpha_{PI3K} = \alpha_{PI3K\_0} \cdot R_0$, resulting in the following equation:

$$PI3K(t) = 0.5 \cdot (1 + k_{PI3K} + 2\alpha_{PI3K\_0} \cdot R_0 \cdot c_2(t) - \sqrt{(1 + k_{PI3K} + 2\alpha_{PI3K\_0} \cdot R_0 \cdot c_2(t))^2 - 8\alpha_{PI3K\_0} \cdot R_0 \cdot c_2(t)})$$

The (2) model also included the equation for 3’IP - the product of PI3K, however the 3’IP kinetics was fast and closely followed PI3K, therefore we skipped this variable for simplicity and to reduce the number of parameters, and included PI3K directly into the equations for PKB activation.

While integrating the PDGF model (2) with the insulin model (1), we kept most of equations from the insulin model, and added the PDGF-related reaction steps into the PKB equations. PDGF-mediated PKB phosphorylation by Tyr was described with the additional terms in the PKB equations with the rate constant *k4d*. Ser phosphorylation of PKB is related to mTORC2 activity, and activation of mTORC2 by active PDGF complex ($c_2$) was described similarly to insulin model, with the rate constant *k5e*. All other steps, including activation of mTORC1 and S6K are described by the insulin model. The parameters *k4d, k5e* were varied to get the same levels of mTORC1a activation (45%) at saturated concentrations of PDGF and insulin (18), while fitting the PKB-Ser timecourses ((2), Fig. S2B). During the fitting the background PKB-Ser473 levels at time=0 (after 600 min without PDGF) were subtracted from each simulated timecourse, as it was done in the experiments. Experimental data on PKB-Ser473 kinetics were scaled with fitting parameter ky_PKB_S473=0.9, giving the best fit.

*Modelling of mTOR activation by NRG.*

Modelling of mTOR activation by neuregulins (NRG) initiated by their binding to ERBB receptors was done similarly to PDGF (6). Only 4 parameters were changed in the PDGF-related part of the model (Table S1). The NRG ligand-receptor binding affinity $K_{D,L}$ was equal to 9 nM (19); the rate constant of the ERBB receptor endocytosis $k_e$ was equal to 0.06 min$^{-1}$ (20). Parameter $k_{PI3K}$ was fitted to timecourses of Ser phosphorylation of PKB after stimulation with various concentrations of NRG (Fig. S2C), with background activities being subtracted from the simulated timecourses, as it was done to the data. Experimental data on PKB-Ser473 kinetics were scaled with fitting parameter ky_PKB_S473=3, giving the best fit. We note that the model failed to describe the initial overshoot observed in the data, possibly attributed to fast, but transient activation of PKB by Erk signalling, which is absent from our model, but potentially contributing to PKB activation by NRG (21).

Table S1. Parameters of the reference model of the mTOR pathway.

| parameter | *k1a* | *k1c* | *k1d* | *k1f* | *k1g* | *k1r* | *k2_0* | *k2a* | *k2c* | *k2b* | *k2d* | *k2f* |
|---|---|---|---|---|---|---|---|---|---|---|---|---|
| value | 0.633 nM$^{-1}$ min$^{-1}$ | 0.877 min$^{-1}$ | 31 min$^{-1}$ | 184000 min$^{-1}$ | 1940 min$^{-1}$ | 0.547 min$^{-1}$ | 0.0423 min$^{-1}$ | 323 min$^{-1}$ | 576000 min$^{-1}$ | 3420 min$^{-1}$ | 281 min$^{-1}$ | 2.91 min$^{-1}$ |
| ref | (1) | (1) | (1) | (1) | (1) | (1) | (1) | (1) | (1) | (1) | (1) | (1) |
| parameter | *k2g* | *k3a* | *k3b* | *k4a* | *k4b* | *k4c* | *k3_0* | *k4f* | *k4h* | *k5* | *k5a1* | *k5b* |
| value | 0.267 | 0.0001 min$^{-1}$ | 0.0988 min$^{-1}$ | 579000 min$^{-1}$ | 34.8 min$^{-1}$ | 446 min$^{-1}$ | 0.01 | 30 min$^{-1}$ | 0.536 min$^{-1}$ | 40 min$^{-1}$ | 0.03 | 24.8 min$^{-1}$ |
| ref | (1) | fit | (1) | (1) | (1) | (1) | fit | fit | (1) | fit | (1) | (1) |
| parameter | *k5d* | *k5c* | *k4_0* | *k4c_0* | *k9f1* | *k9b1* | *k9f2* | *k9b2* | *km9* | *kiPTEN* | *kiTSC* | |
| value | 1.06 min$^{-1}$ | 2 min$^{-1}$ | 4 min$^{-1}$ | 0.3 min$^{-1}$ | 0.9 min$^{-1}$ | 0.0444 min$^{-1}$ | 333 min$^{-1}$ | 31 min$^{-1}$ | 58.7 | $1\cdot10^{-5}$ | 0.001 | |
| ref | (1) | fit | fit | fit | fit | (1) | (1) | (1) | (1) | (12) | fit | |
| parameter | $K_{D,L}$ | $k_f$ | $k_x$ | $R_0$ | $k_e$ | $k_{L,max}$ | $K_{M,L}$ | $\alpha_{PI3K\_0}$ (PDGF) | $k_{PI3K}$ (PDGF) | *k4d* | *k5e* | $k_{-x}$ |
| value | 1.5 nM (pdgf) 9 nM (nrg) | 1 nM$^{-1}$ min$^{-1}$ | 30 min$^{-1}$ | 0.01 | 0.2 min$^{-1}$ (pdgf); 0.06 min$^{-1}$ (nrg) | 0.011 min$^{-1}$ | 1.66 nM | 8000 | 0.3 (pdgf) 100 (nrg) | 30 min$^{-1}$ | 0.2 min$^{-1}$ | 0.07 min$^{-1}$ |
| ref | (2, 19) | (17) | (2) | fit | (2, 20) | (2) | (2) | (2)* | (2), fit | fit | fit | (2) |

* $\alpha_{PI3K\_0}$ is equal to $\alpha_{PI3K}/R_0$ = 80/0.01, where $\alpha_{PI3K}$ value is obtained from (2); PTEN=TSC=1

#### ***1.5. Selection of GFs to be included in the model***

In our brain-specific model we used publicly available snRNA-seq data on human brain samples from cortex and hippocampus of 8 healthy donors (control) and 9 AD donors (22). The data were downloaded from https://twc-stanford.shinyapps.io/human_bbb/ and were represented by gene expression averaged between the samples, for each cell type. We collated the expression of mTOR pathway genes (below) and firstly determined which GF receptors are expressed in the brain cells together with their cognate GFs, activating canonical mTOR pathway. The GFs were analysed by their group expression – the total expression of all GFs from the same family (Fig. S3). The expression of GFs of the EGF family was high and mainly represented by neuregulins (NRG), with their ERBB receptors being highly expressed in several types of brain cells (Fig. S3). The high-affinity PDGF receptors were expressed in pericytes, smooth muscle cells (SMC), oligodendrocyte precursor cells (OPC) and neurons, and PDGF GFs were widely expressed (Fig. S3). Insulin receptors expressed in most of brain cells (Fig. S3). FGF was moderately expressed, but its activity via the canonical mTOR pathway in adult brain is mainly related to stress conditions (survival, apoptosis) (23, 24), while the role of FGF in proliferation is mainly mediated by MAPK signalling (25, 26). VEGF receptors were mainly represented by the inhibitory and pathologically involved FLT receptor (27), while the expression of the mTOR-activating KDF receptor was very low (Fig. S3). Taking all expression and mTOR-activation characteristics into account we concluded that IGFs, NRGs and PDGFs represent the main groups of GFs expressed in mature brain cells activating the canonical mTOR pathway under normal conditions.

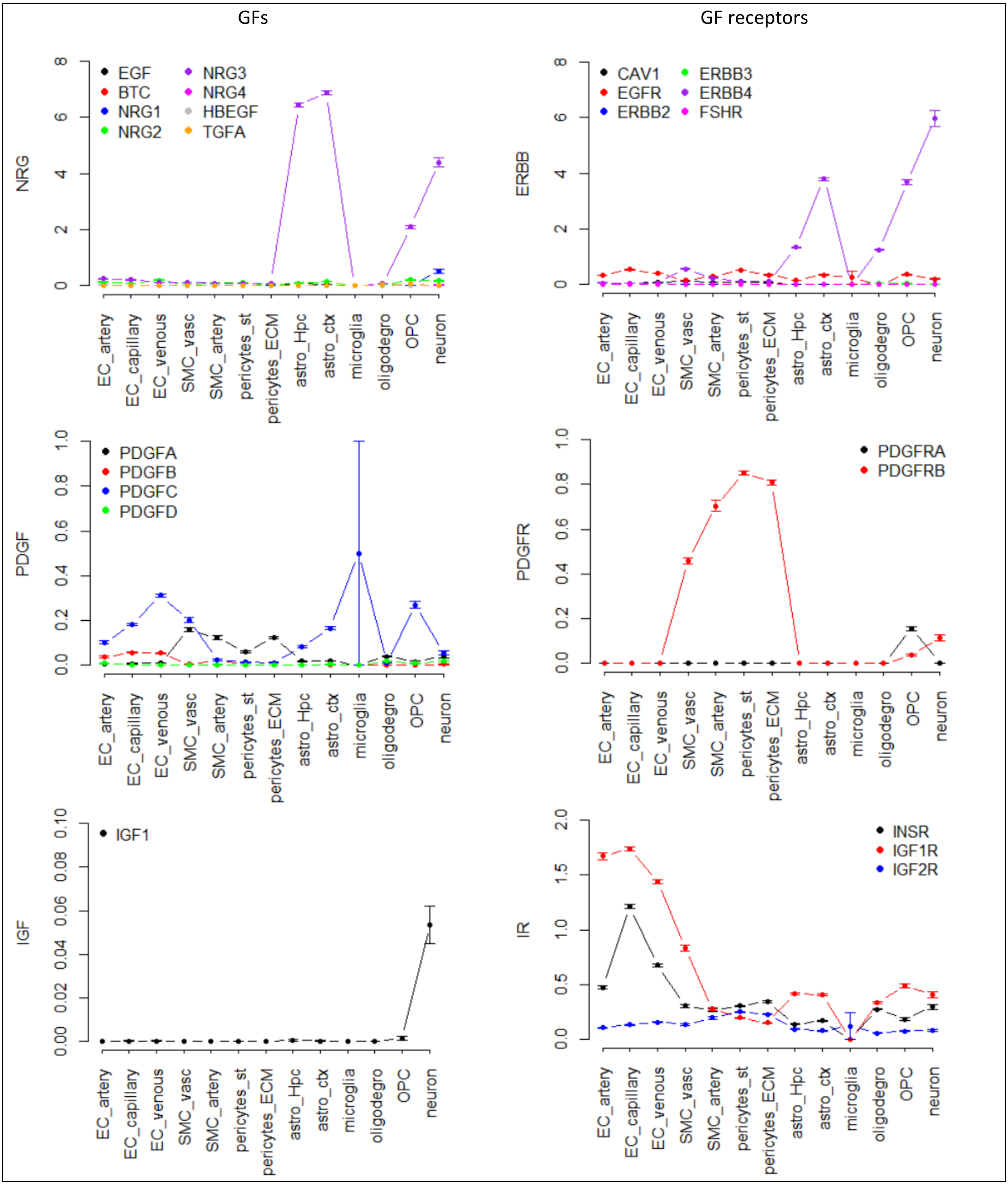

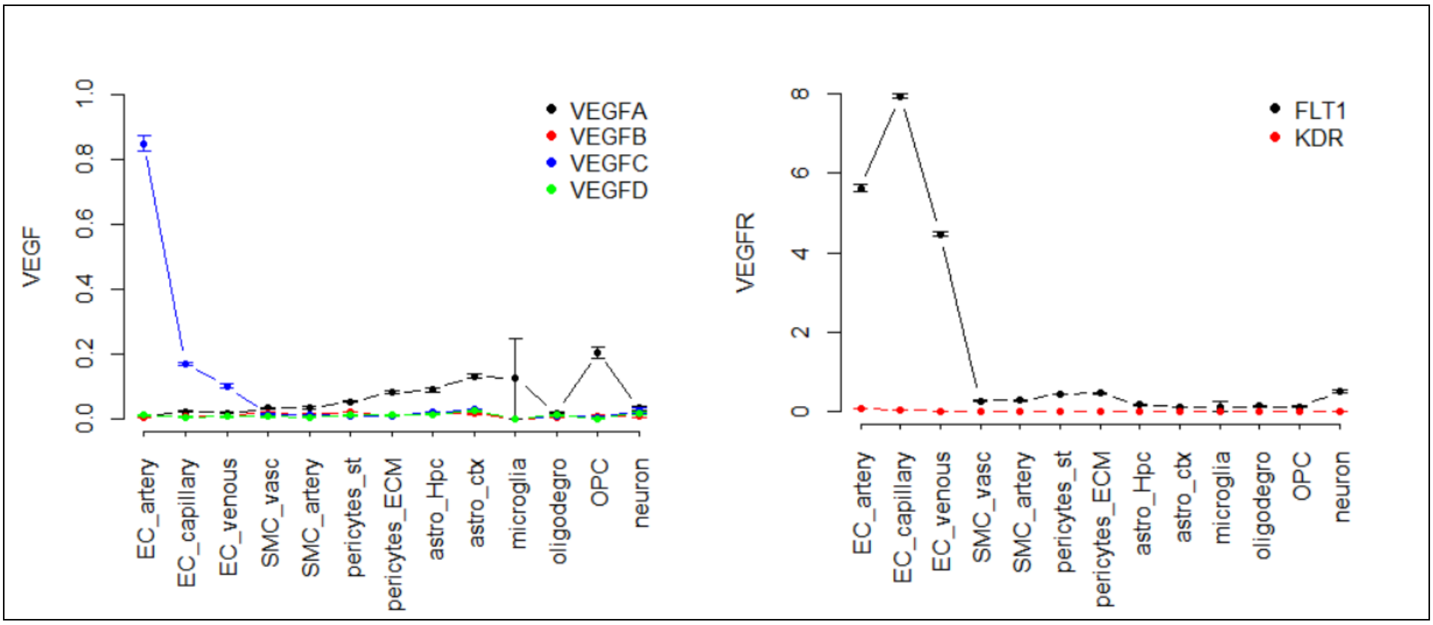


Figure S3. Expression of GFs and GF receptors participating in canonical mTOR pathway activation in brain cells. The expression of genes in each group of GFs and their receptors is shown for the different cell types (22). The expression is averaged between cortex and hippocampus samples from 8 healthy donors. Error bars show SEM. The names of GF groups are shown in y axis. Individual gene names are shown in the legends. Sub-population designations are: pericytes_st (solute transport pericytes) and pericytes_ECM (ECM-regulating pericytes); astro-Hpc (hippocampal astrocytes) and astro-ctx (cortical astrocytes).

#### ***1.6. Simulations of random fluctuations in the abundance of mTOR components***

Random fluctuations in the abundances of mTOR model components were simulated by sampling the abundances from log-normal distribution (with mean and SD equal to 1), providing a good approximation of gene expression profiles (28). Since our aim was to compare the effects of random fluctuations with the effects of intercellular variations in gene expression in the brain-specific model (described below), we perturbed only those variables, which were varied in the brain-specific model (IR, PDGFR, NRGR, IRS1, PTEN, PKB, and S6K). 1000 “random cells” were generated, each having random variations in these variable, and S6Ka maximum after 2h of mTOR stimulation with 10 nM of GFs were calculated. Changes in the abundances of individual components had different effects of the S6Ka (Fig. S4).

To quantify the combined effect of variations in the abundances of multiple variables, we defined the functions of various combinations of abundances. The functions included multiplications for abundances of mTOR-activating variables and divisions for inhibitory variables (PTEN and TSC). Various combinations of random fluctuations in abundances were tested (Fig. S5 A) and compared to the brain-specific variations (Fig. S5 B). Microglia cell type was removed from this analysis due to low and therefore noisy expression of mTOR components. Only cells expressing GF receptors at levels ≥ 1% from maximum were used in the stimulations (all cells for insulin and NRG, but some cell types with negligible PDGFR were omitted from PDGF stimulations, Fig. S5).

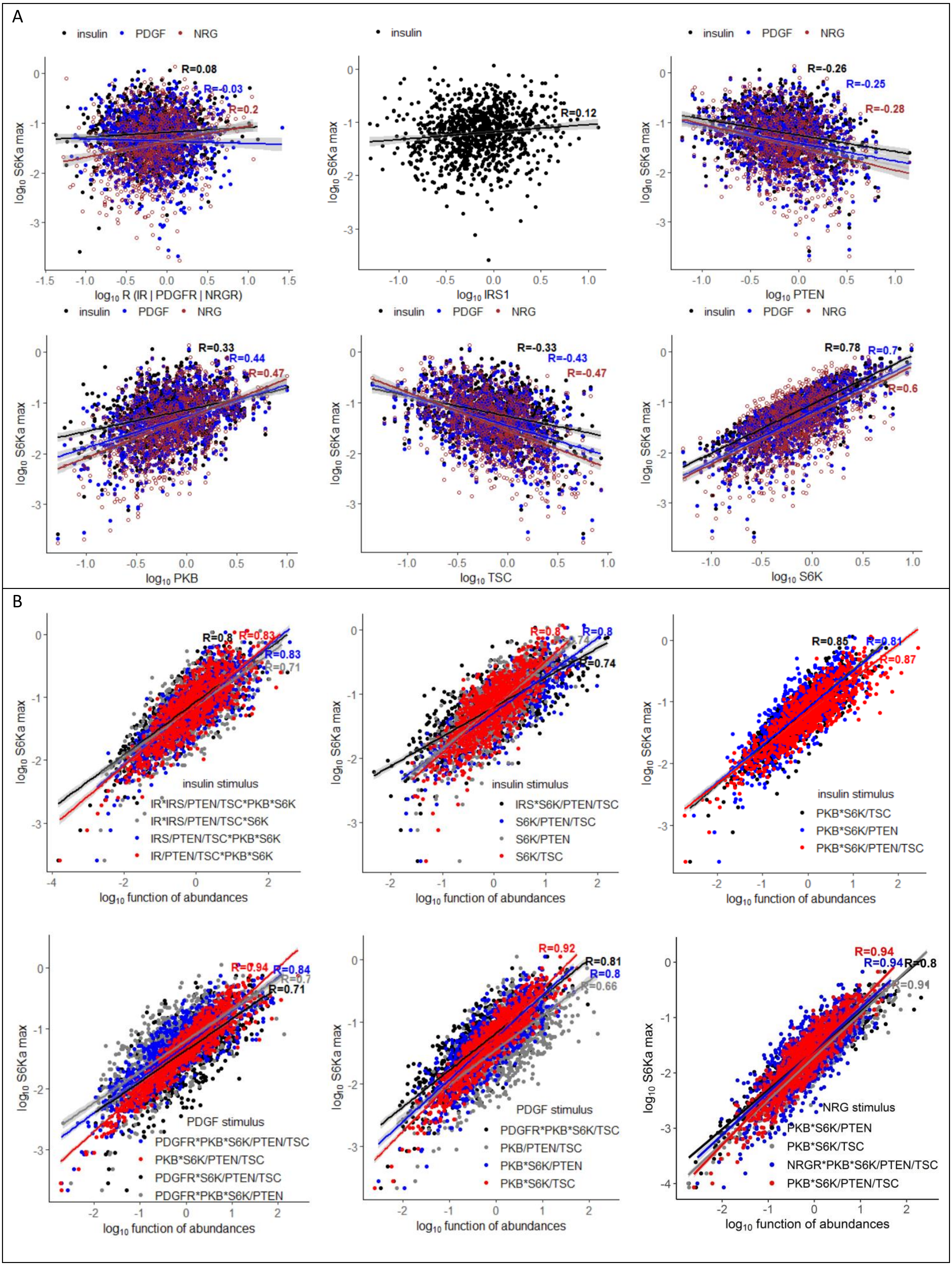


Figure S4. Effect of random fluctuations in abundance of mTOR components on the maximum S6Ka. Calculations of S6Ka were done with the reference model after 2h of stimulation with 10 nM of GFs, with varying abundances of mTOR components. Plots shows linear correlation between $\log_{10}$ of S6Ka and either single variable abundances (A), or multiple variable abundances (legends on B). The effect of IRS1 was analysed only for insulin stimulation. The effects of GF receptors were analysed for their respective stimulus: IR for insulin, PDGFR for PDGF and NRGR/ERBB for NRG.

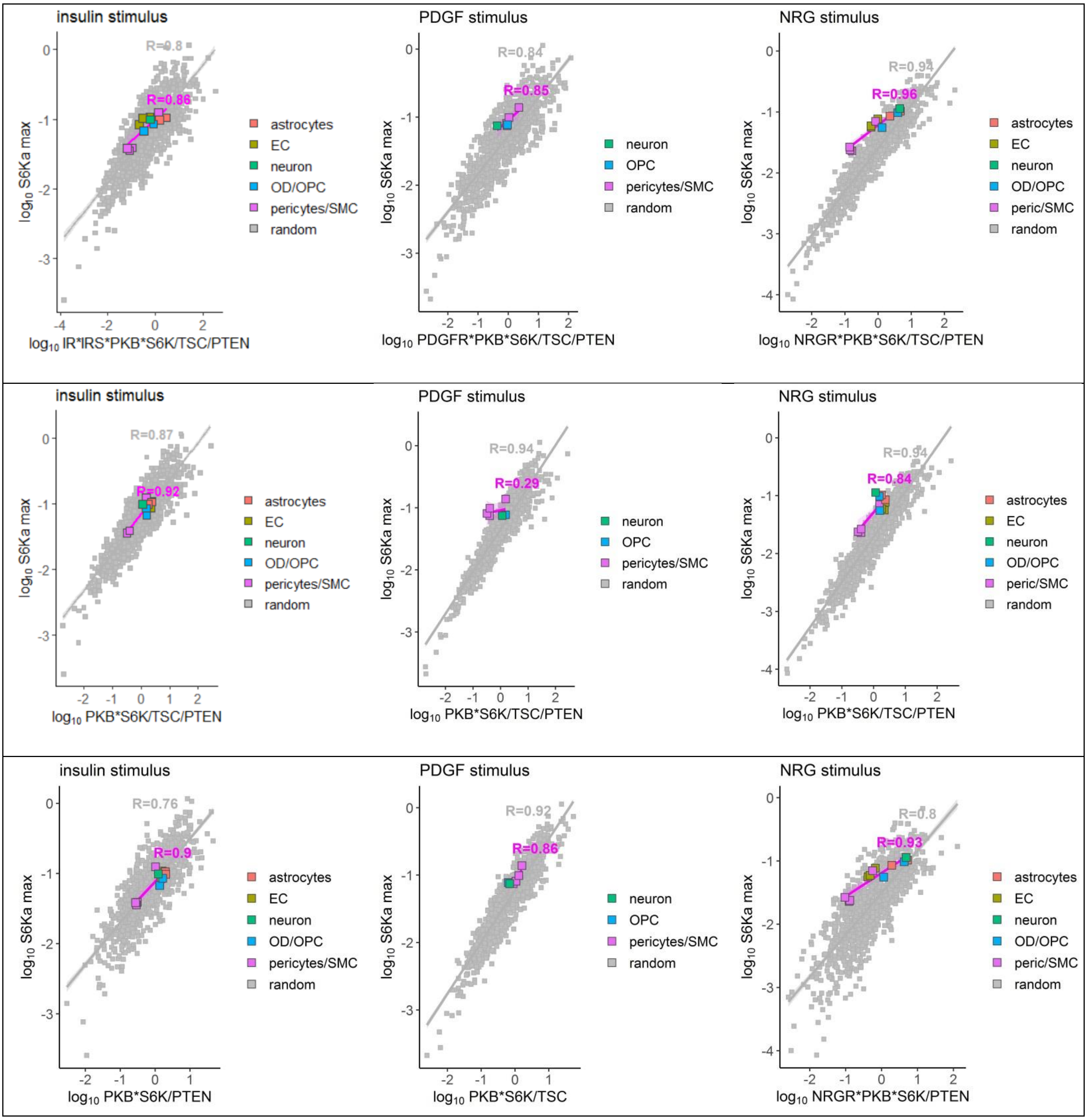


Figure S5. Comparison of random variations in the abundance of mTOR components with brain cell-specific abundance profiles. S6Ka was calculated after 2h of stimulation with 10 nM of GFs. The linear correlation between $\log_{10}$ of S6Ka and various functions of abundances (x axis) for simulated “random cells” (with random fluctuations in abundances, grey) and the brain cells (coloured). Different types of EC (artery, capillary and venous), pericytes (solute transporting and ECM-regulating), SMC (vascular, artery) and astrocytes (hippocampal and cortex) are represented by the same colours for simplicity.

### 2. Modelling the mTOR pathway in brain cells

After building the reference mTOR model, we used the public snRNA-seq data (22) to infer the relative abundance of mTOR pathway components in different types of brain cells. In the brain-specific model, the total concentration of a model component was represented by normalized sum of expression of related genes (Fig. S6, Table S2). The expression was averaged between the brain samples of 8 healthy donors or 9 AD patients. The insulin model was developed using adipocyte cell cultures (1), and our preliminary comparison of mTOR gene expression between the brain cells (22) and adipocytes (29) suggested that expression was higher in adipocytes. Therefore, we used the upper quantile normalization between the cell types with non-zero summary expression in each group of genes, corresponding to total concentrations of IR, IRS, PKB, PTEN, TSC, S6K, PDGFR and NRGR. Zero summary expression below the snRNA-seq detection limit was replaced by half of minimal expression (amongst all samples) in each group. We note that the expression of some mTOR components was highly variable in microglia (Fig. S6), possibly due to variability in functional states of microglia cells and/or relatively low number of microglia cells obtained from the vascular tissues (30). Therefore, the model describes average properties of mTOR in microglia population. The expression of genes coding components of mTORC1 and mTORC2 complexes, such as MTOR and RPTOR was relatively constant in (22) dataset. The regulation of mTORC1 and mTORC2 is complex, incompletely understood, and includes post-translational modifications of multiple proteins with currently unknown kinetics. Therefore, the amounts of mTORC1 and mTORC2 were kept equal to 1, similar to the reference model. The initial concentrations of non-active/non-phosphorylated components were equal to their total concentrations and the initial concentrations of active/phosphorylated components were equal to zero. All simulations were initially run for 600 min without GFs, followed by step-wise increase in GF concentrations.

Sensitivity of S6Ka to changes in the levels of PTEN or TSC was defined as fold change in S6Ka upon 4-fold changes in PTEN or TSC (2-fold decrease relative to 2-fold increase). S6Ka was calculated after 2 h of mTOR stimulation with the GF concentration providing maximal activation (in the range of GFs < 10 nM).

mTOR activity was characterized by maximal S6K activity and the effective ligand dose (EC50) (or potency - the reciprocal EC50). Both S6Ka and potency were lower for NRG stimulation compared to insulin and PDGF, therefore we scaled them on Figs. 3,4 and Fig. S8 for easy comparison between stimuli. For NRG stimulation we multiplied S6Ka to 2 and potency to 5.

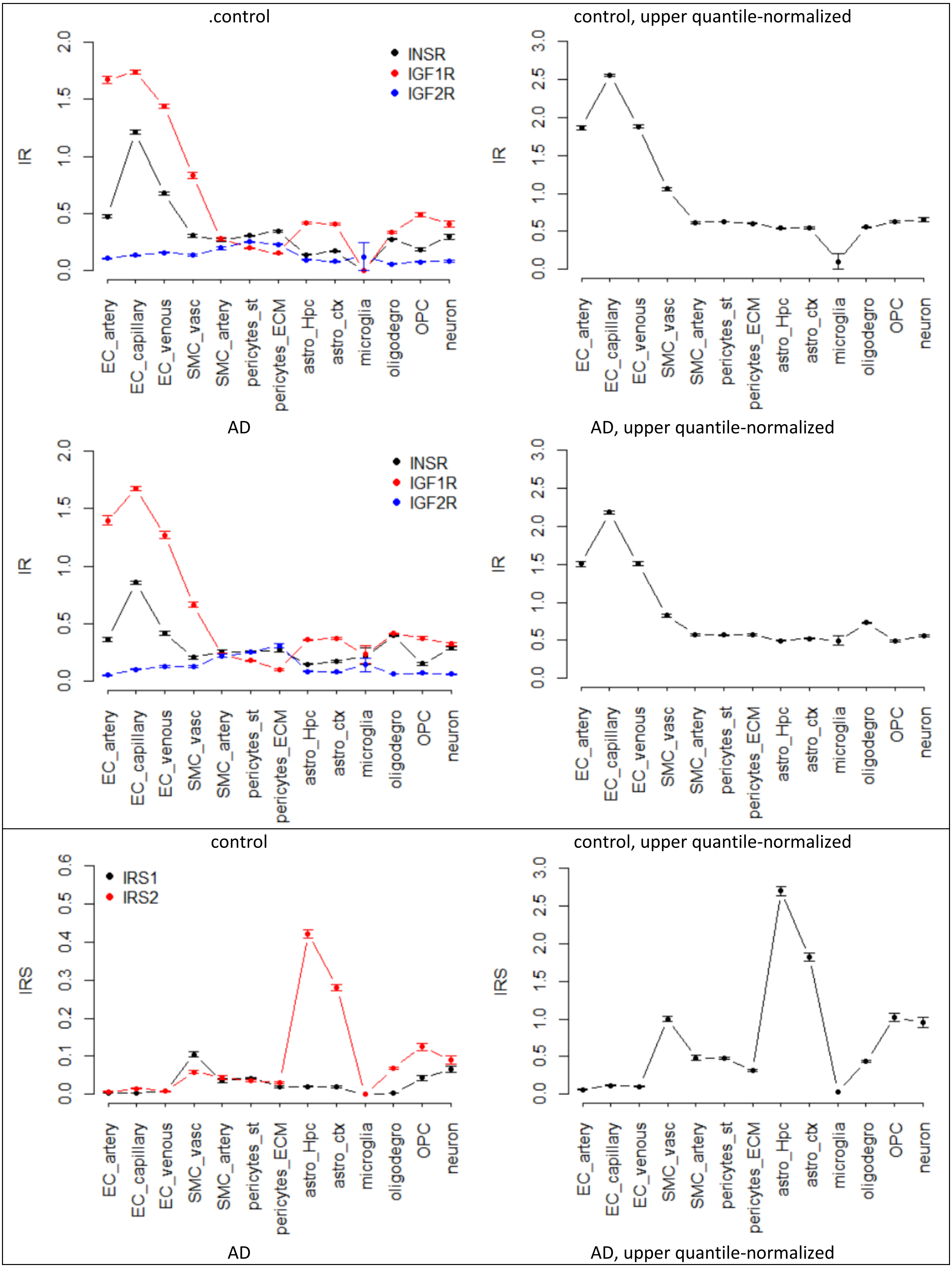
.control
control, upper quantile-normalized
AD
AD, upper quantile-normalized
control
control, upper quantile-normalized
AD
AD, upper quantile-normalized
INSR
IGF1R
IGF2R
IRS1
IRS2
IR
IRS
0.0
0.5
1.0
1.5
2.0
2.5
3.0
0.1
0.2
0.3
0.4
0.5
0.6
EC_artery
EC_capillary
EC_venous
SMC_vasc
SMC_artery
pericytes_st
pericytes_ECM
astro_Hpc
astro_ctx
microglia
oligodegro
OPC
neuron

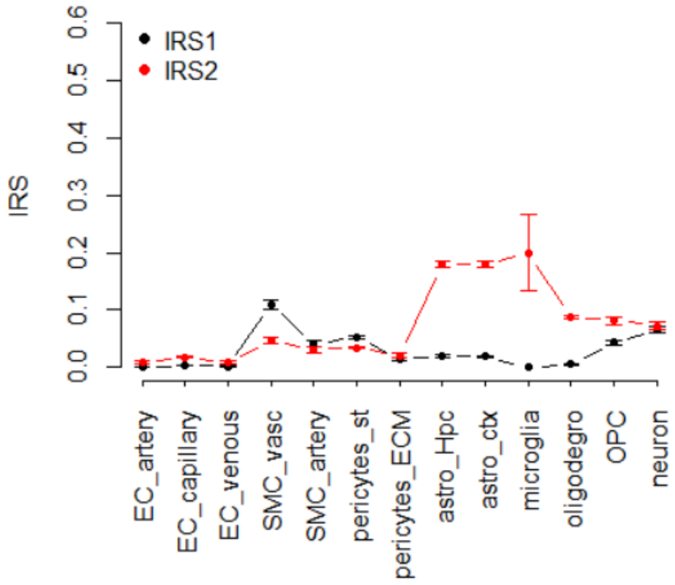


control

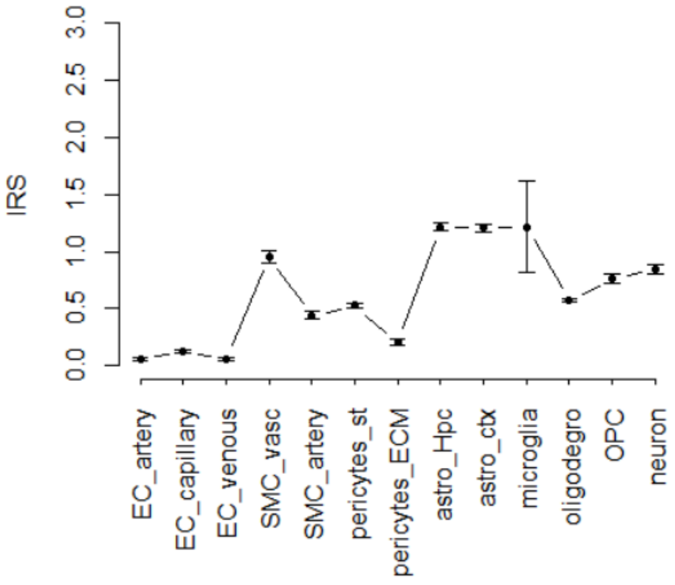


control, upper quantile-normalized

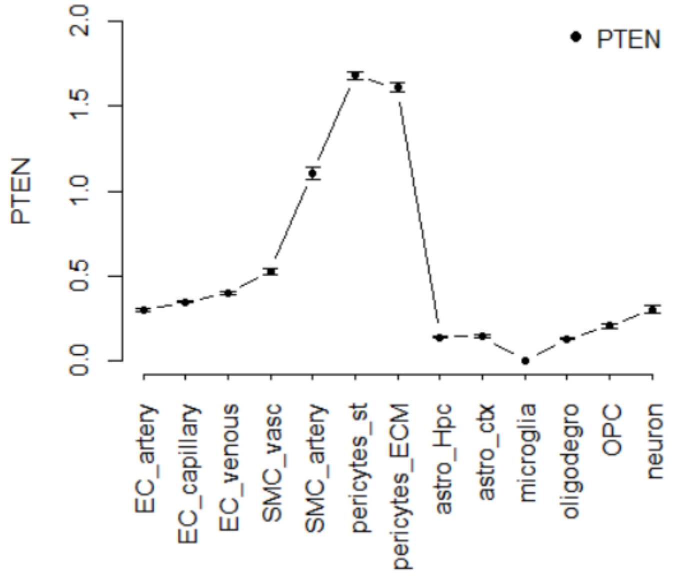


AD

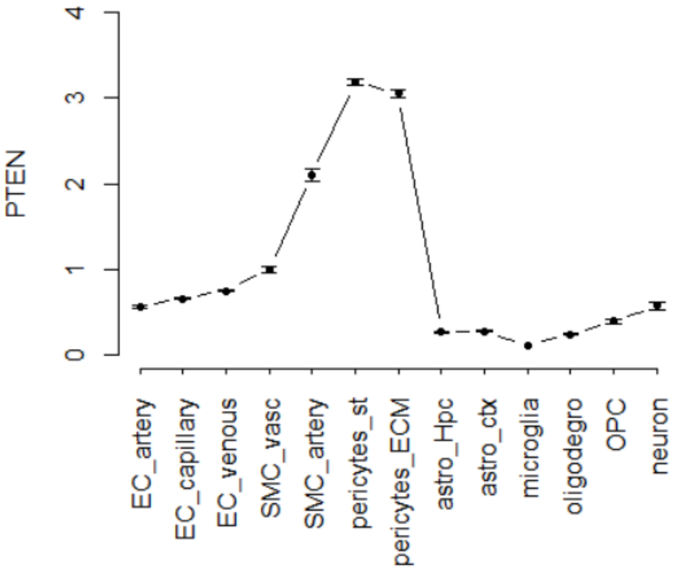


AD, upper quantile-normalized

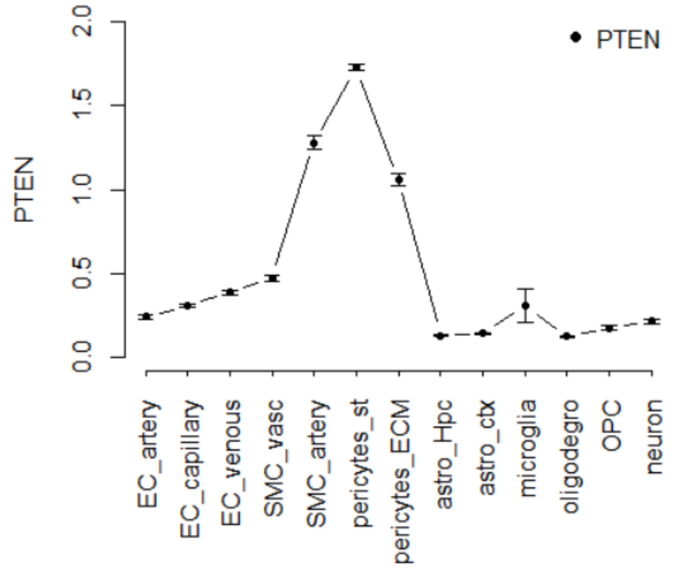


control

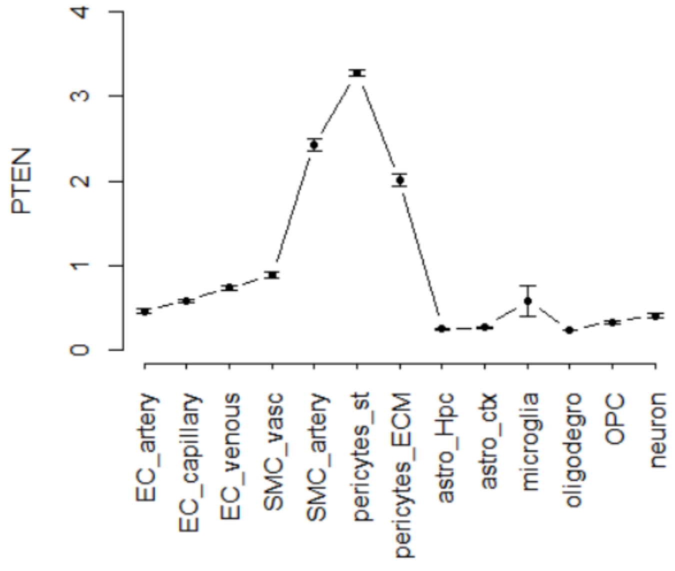


control, upper quantile-normalized

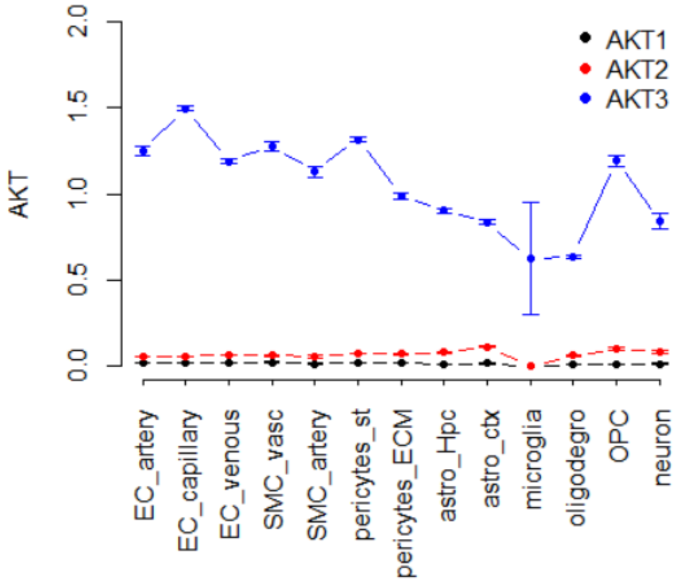


AD

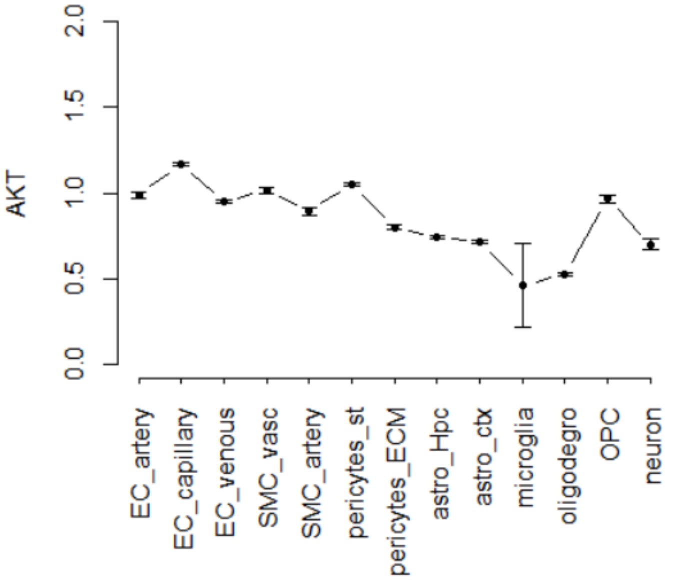


AD, upper quantile-normalized

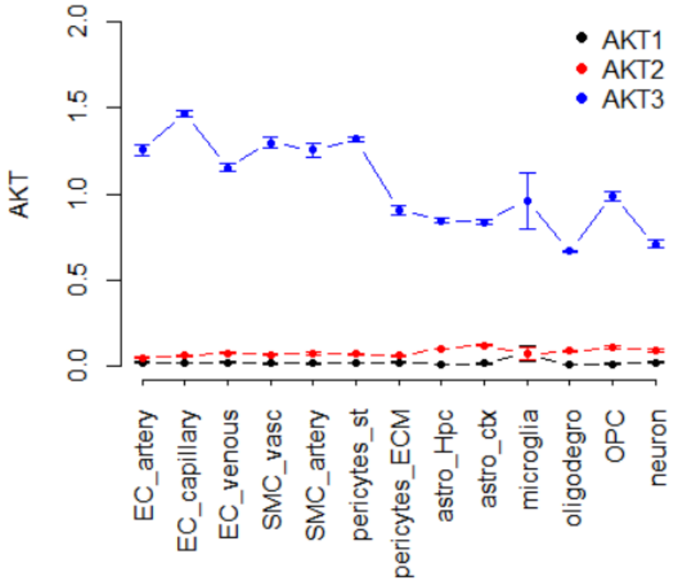


control

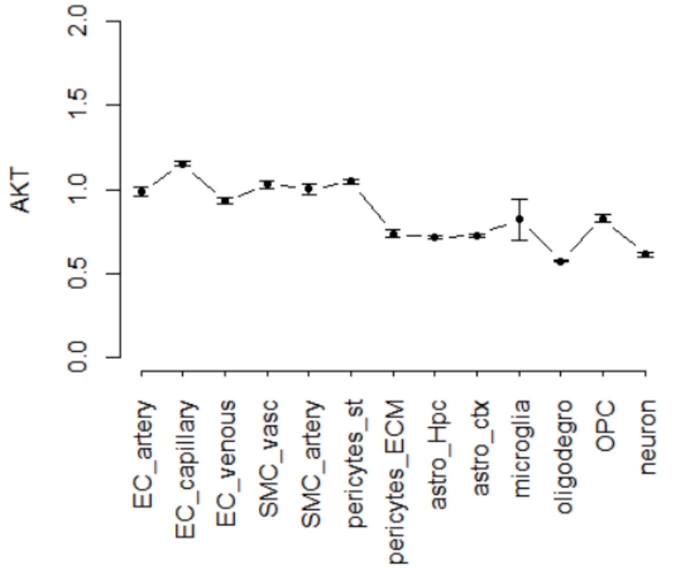


control, upper quantile-normalized

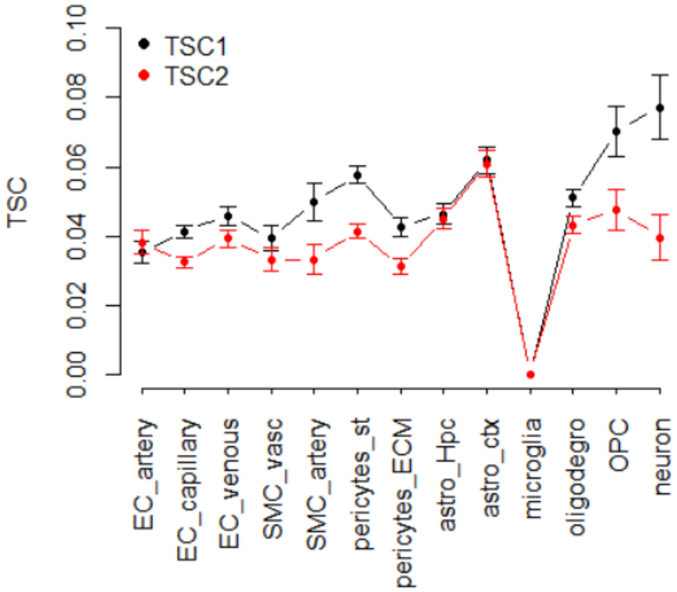


AD

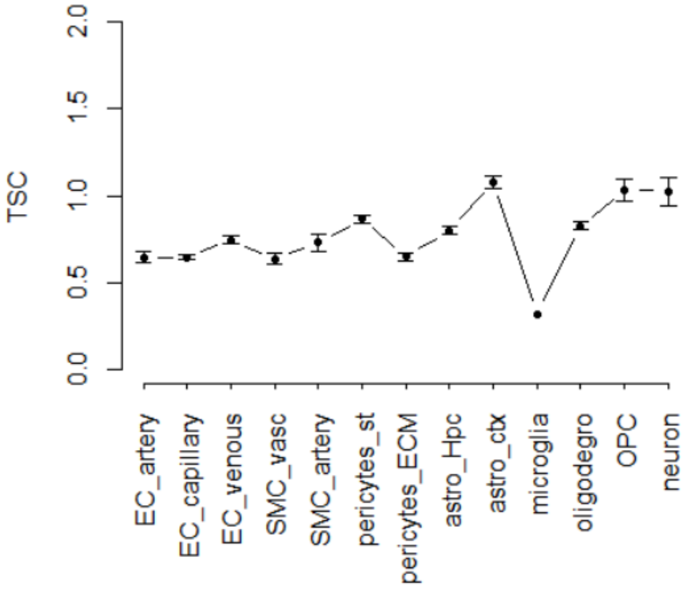


AD, upper quantile-normalized

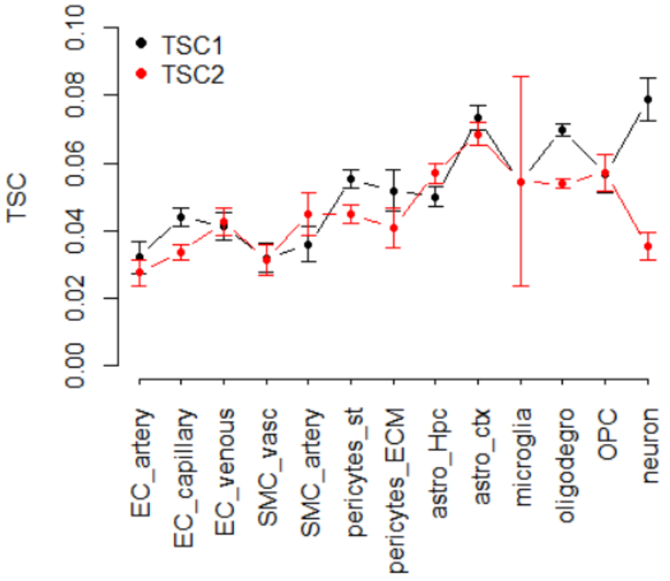


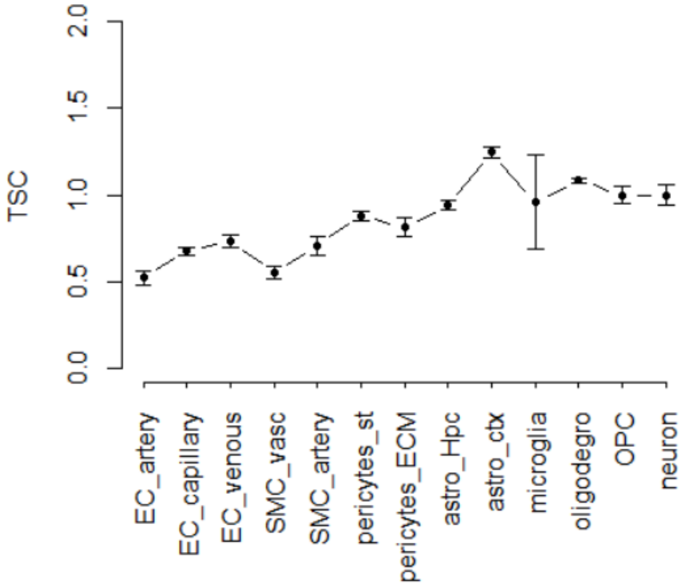


control

control, upper quantile-normalized

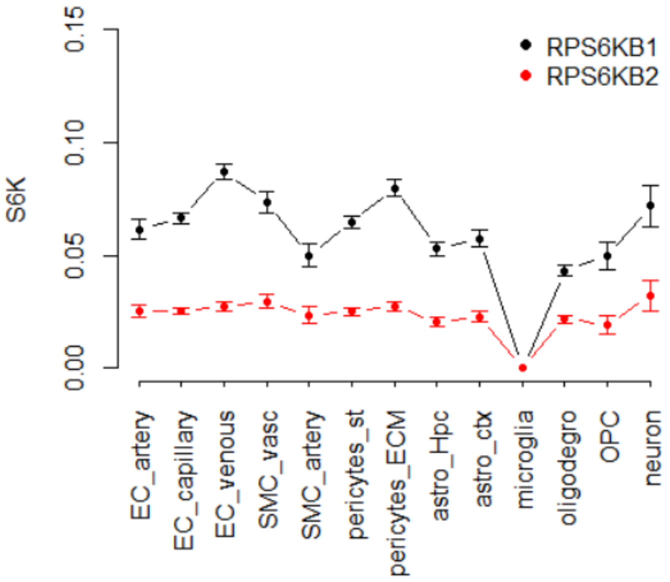


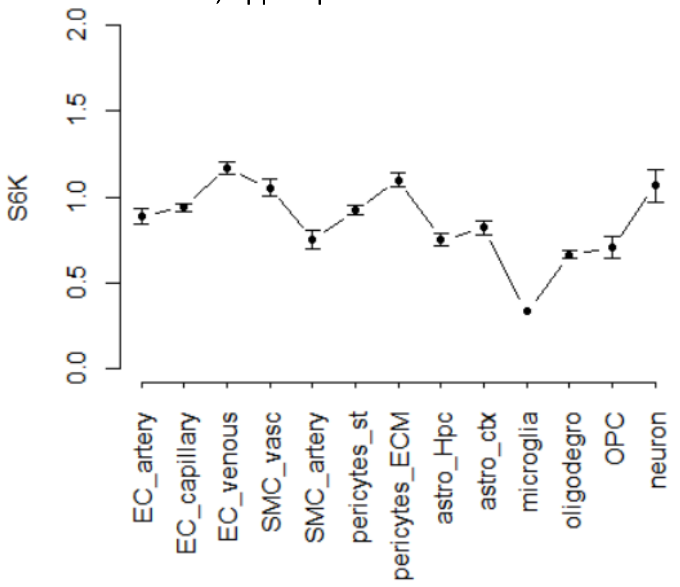


AD

AD, upper quantile-normalized

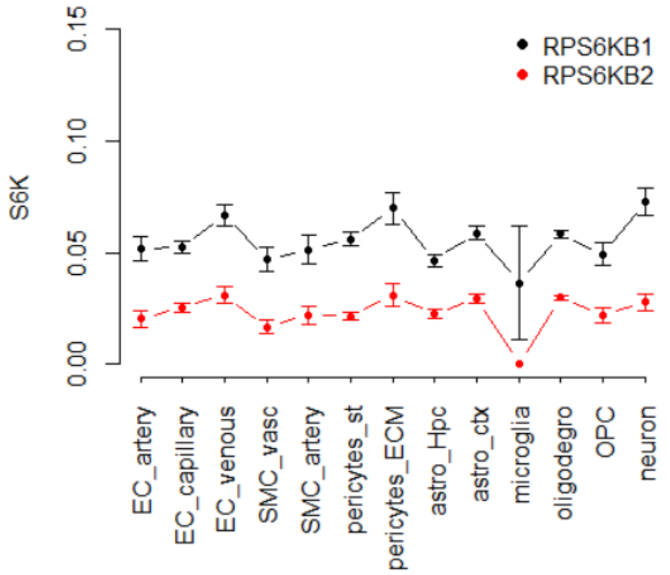


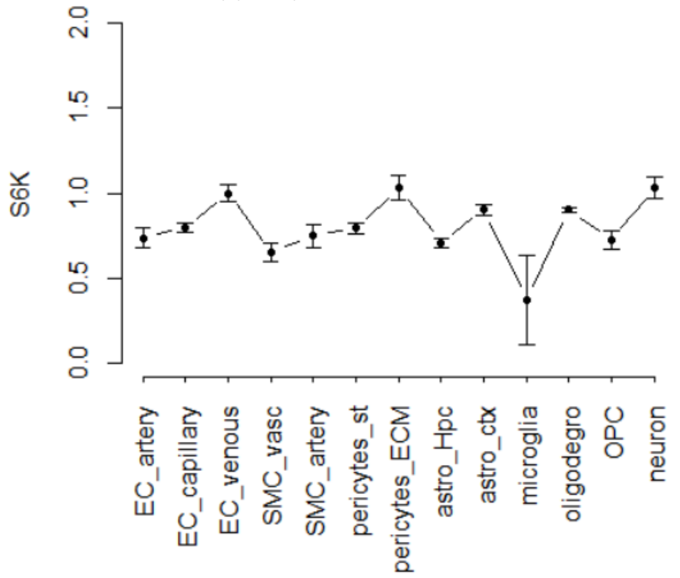


control

control, upper quantile-normalized

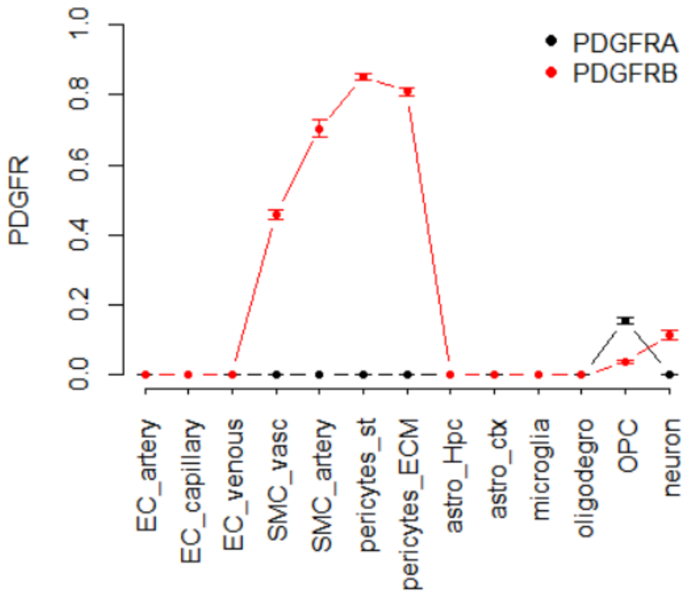


AD

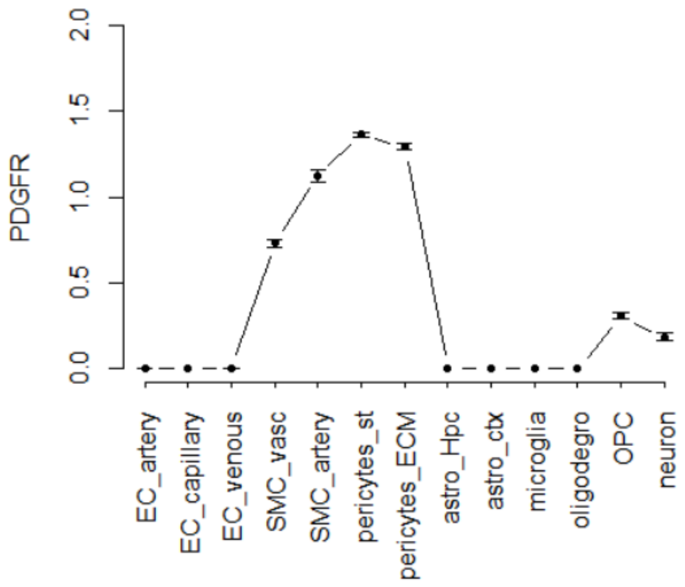


AD, upper quantile-normalized

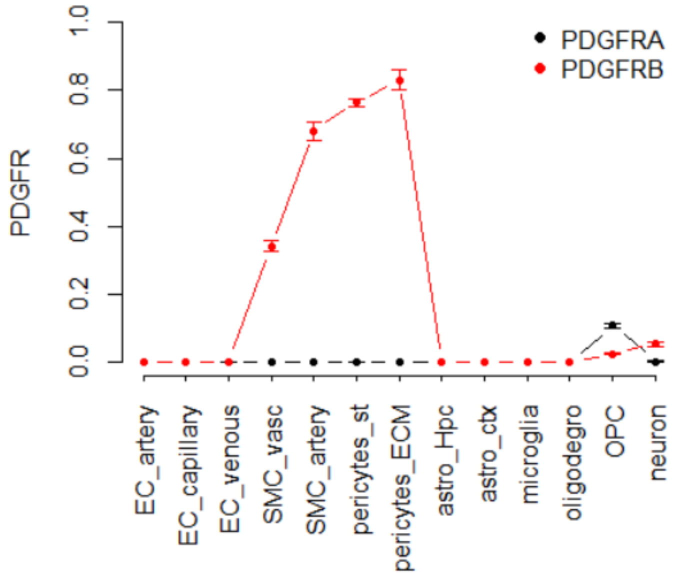


control

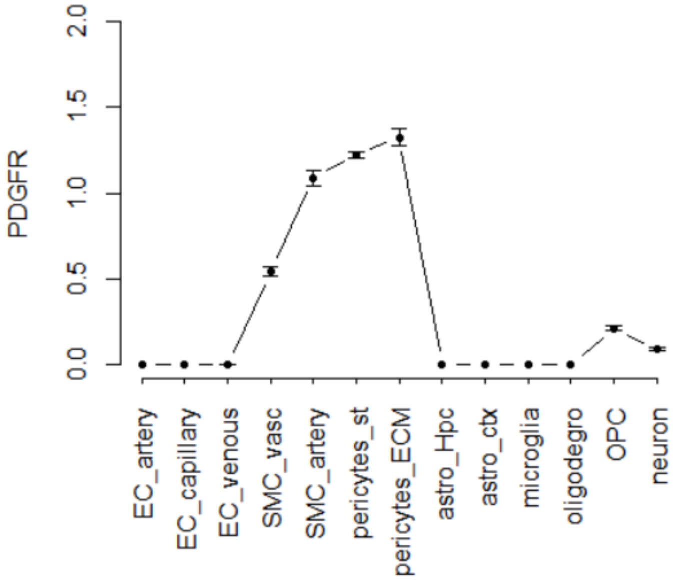


control, upper quantile-normalized

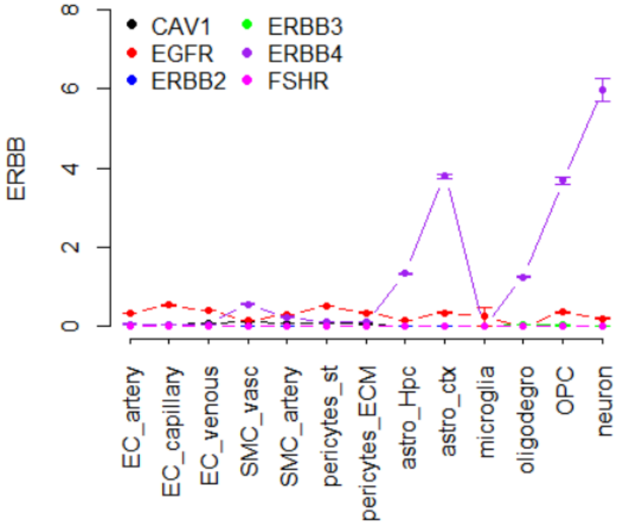


AD

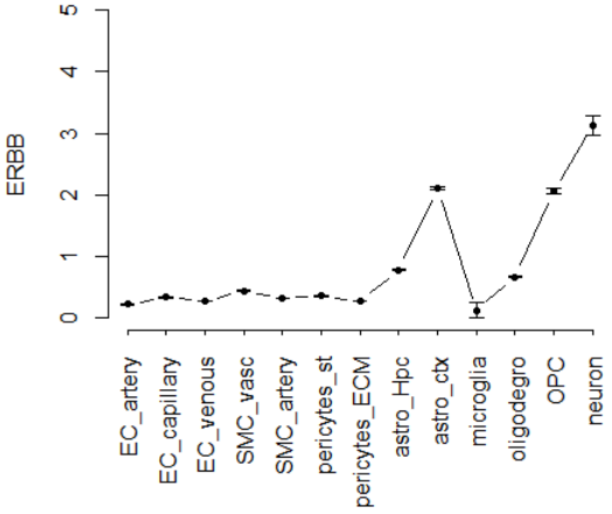


AD, upper quantile-normalized

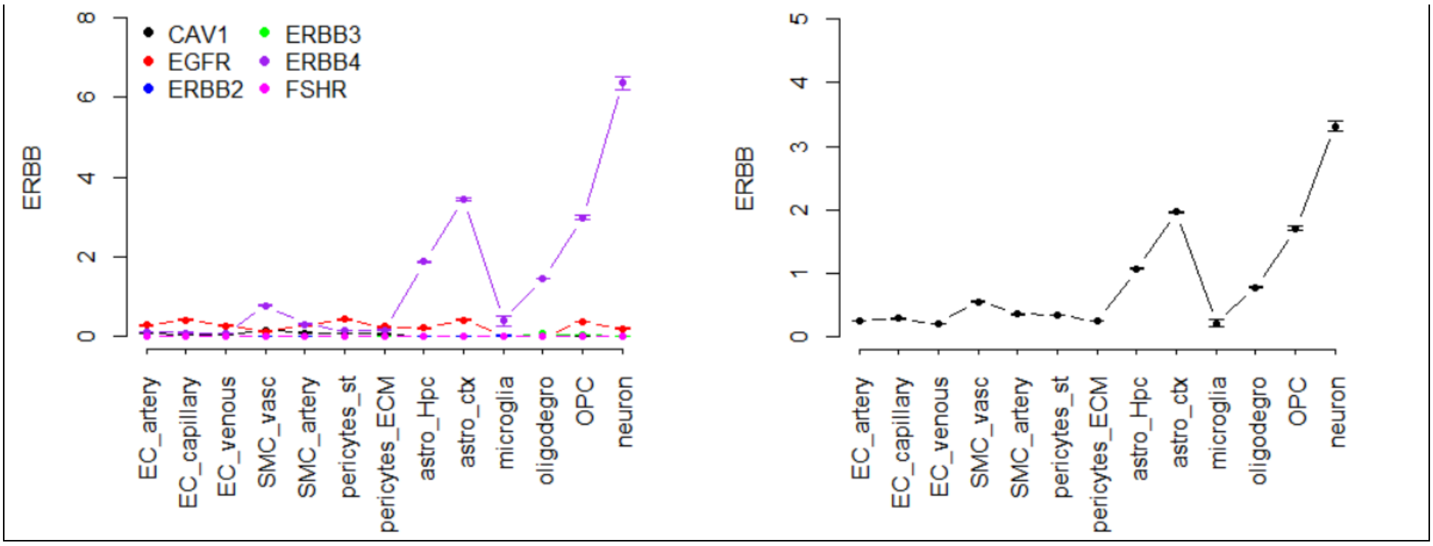


Figure S6. Expression of mTOR pathway components in brain cells. Left: Expression of genes corresponding to each mTOR component in the model, for different cell types. The expression is averaged between the samples (control or AD) (22). Right: Upper quantile normalized sum of expression in each gene group used as an input into the model for the total amount of the respective mTOR component.

Table S2. Normalized expression of brain cell model components for healthy conditions. Expression was summarized between genes in each group corresponding to a model variable and the total expression was upper quantile normalized.

| gene | EC artery | EC capillary | EC venous | SMC vascul | SMC artery | Pericyte st | Pericyte ECM | Astro Hpc | Astro Ctx | microgl | OD | OPC | neuron |
|---|---|---|---|---|---|---|---|---|---|---|---|---|---|
| PDGFR | 0.005 | 0.001 | 0.003 | 1.48 | 2.27 | 2.76 | 2.61 | $5\cdot10^{-4}$ | $5\cdot10^{-4}$ | $5\cdot10^{-4}$ | 0.002 | 0.62 | 0.37 |
| ERBB | 0.28 | 0.42 | 0.34 | 0.55 | 0.41 | 0.45 | 0.34 | 0.97 | 2.64 | 0.16 | 0.84 | 2.58 | 3.91 |
| IR | 1.79 | 2.46 | 1.81 | 1.02 | 0.60 | 0.60 | 0.58 | 0.52 | 0.53 | 0.10 | 0.54 | 0.60 | 0.63 |
| IRS | 0.06 | 0.11 | 0.10 | 0.90 | 0.43 | 0.43 | 0.28 | 2.43 | 1.64 | 0.03 | 0.39 | 0.92 | 0.86 |
| PTEN | 0.60 | 0.69 | 0.79 | 1.05 | 2.20 | 3.34 | 3.21 | 0.28 | 0.29 | 0.13 | 0.26 | 0.41 | 0.61 |
| PKB | 1.00 | 1.19 | 0.96 | 1.03 | 0.91 | 1.06 | 0.81 | 0.75 | 0.73 | 0.47 | 0.53 | 0.98 | 0.71 |
| TSC | 0.68 | 0.68 | 0.78 | 0.67 | 0.76 | 0.91 | 0.68 | 0.84 | 1.13 | 0.32 | 0.87 | 1.08 | 1.07 |
| S6K | 0.88 | 0.93 | 1.15 | 1.04 | 0.74 | 0.91 | 1.08 | 0.74 | 0.81 | 0.13 | 0.66 | 0.70 | 1.05 |

The R code of the model is freely available at https://github.com/alex297/mTOR
